# Neuromorphic hierarchical modular reservoirs

**DOI:** 10.1101/2025.06.20.660760

**Authors:** Filip Milisav, Andrea I. Luppi, Laura E. Suárez, Guillaume Lajoie, Bratislav Misic

## Abstract

Modularity is a fundamental principle of brain organization, reflected in the presence of segregated sub-networks that enable specialized information processing. These small, densely connected modules are often nested within larger, higher-order modules, giving rise to a hierarchical modular architecture. This structure is posited to balance information segregation in specialized neuronal communities and global integration via intermodular communication. Yet, how hierarchical modularity shapes network function remains unclear. Here we introduce a simple blockmodeling framework for generating and comparing multi-level hierarchical modular networks and implement them as recurrent neural network reservoirs to evaluate their computational capacity. We show that hierarchical modular networks enhance memory capacity, support multitasking, and give rise to a broader range of temporal dynamics compared to strictly modular and random networks. These functional advantages can be traced to topological features enriched in hierarchical modular networks, which include reciprocal and cyclic network motifs. To test whether the computational advantages of hierarchical modularity subsist in empirical human brain structural connectivity patterns, we develop a novel hierarchical modularity-preserving network null model, allowing us to isolate the positive effect of empirical hierarchical modularity patterns on memory capacity. To evaluate the biomimetic validity of connectome-informed reservoir dynamics, we compare reservoir timescales to empirical brain timescales derived from MEG data and find that hierarchical modularity contributes to shaping brain-like neural timescales. Altogether, across multiple benchmarks, these results show that hierarchical modularity endows networks with computationally advantageous properties, providing insight into the relationship between neural network structure and function with potential applications for the design of neuromorphic computing architectures.

## INTRODUCTION

Modularity is a fundamental principle in neuroscience, shaping our understanding of neural architecture, dynamics, and function. The brain is not a homogeneous system, but a complex network, composed of distinct yet interacting modules. In network neuroscience, modularity is a defining feature of brain organization, manifesting in the segregation of structural and functional subnetworks that support specialized processing [28, 75, 114, 170]. Beyond neurobiological organization, modularity is also increasingly studied in artificial intelligence [174], both as a network feature [2, 31, 36, 56, 68, 94, 99, 107, 177] and as a design principle [3, 5, 26, 67, 92, 119, 140, 141, 148]. In this context, a module can consist of any neural architecture component, from a layer to a whole network as part of an ensemble [55, 81, 140, 174]. Modular systems leverage intrinsic or imposed modularity in data and tasks via routing functions to select relevant modules and aggregation functions to combine their outputs [140, 174]. Such approaches have been shown to offer key advantages such as enhanced interpretability [92, 99, 141], improved efficiency and scalability [5, 55, 119, 140, 159], better generalization and knowledge transfer [5, 25, 26, 92, 119, 141, 156], increased robustness [25], and ease of knowledge retention [48], enabling systems to decompose complex tasks and recombine learned solutions [5, 26, 92, 99, 141, 148, 156].

These notions echo a deeply rooted principle in cognitive science, which suggests that cognitive processes operate within informationally encapsulated domains [33, 57], highlighting modularity as a key characteristic of mental architecture. More recent perspectives suggest that cognition emerges from a hierarchy of increasingly polyfunctional nested circuits [15, 76, 114, 115, 135], dynamically recruited to support both domain-specific and integrative processes [4, 162]. Thus, understanding modularity is essential not only for characterizing the brain’s structural and functional topology but also for elucidating the computational principles that underlie human cognition.

Small densely connected modules can themselves be embedded into higher-order modules, leading to hierarchical modular networks. Such a multi-scale organization has been hypothesized to maintain a balance between information segregation in specialized communities and global integration via intermodular communication [76, 114]. Previous work has focused on characterizing the dynamical effects of hierarchical modularity in synthetic neural networks [21, 125, 186]. Notably, using both simple spreading models and spiking neural networks, it was found that hierarchical modularity supports criticality [87, 121, 152, 190, 191], a dynamical regime characterized by advantageous computational properties [13, 20, 32, 42, 51, 63, 91, 95, 97, 131, 160, 161]. Hierarchical modularity has also been associated with increased functional diversity [189, 198], but only as derived from functional connectivity. Yet, how hierarchical modularity shapes cognitive function remains unclear.

Here, we take advantage of reservoir computing [45, 83, 105], a machine learning framework particularly well suited to the implementation of neuromorphic networks [172]. This allows us to move away from the common operationalization of function as statistical associations between neural activity time-courses to instead re-conceptualize it as a computational property in the context of cognitive tasks [104, 172, 173]. In reservoir computing, the classic architecture consists of a recurrent neural network (RNN) of nonlinear neurons complemented by a linear readout module [37, 83, 103]. Only the readout module is trained, allowing us to constrain the network with arbitrary connectivity patterns that remain unchanged throughout learning. Furthermore, since the dynamics of the reservoir are constrained by its fixed wiring, they can be tuned and maintained by the experimenter [173]. This allows the exploration of the interplay between structure, dynamics, and cognitive function.

While a small number of network properties have been related to performance in reservoir computing tasks [1, 22, 38, 44, 85, 90, 112, 113, 147, 195], how neural network architecture supports cognitive capacity remains largely unknown. Recent evidence indicates that an optimal level of modularity can enhance reservoir memory [147] and multitasking capacity [102] by balancing information segregation and integration. Here, we propose that hierarchical modularity can more robustly strike this balance, leading to improved cognitive capacity (Fig. 1). To test this hypothesis, we generate synthetic multi-level hierarchical modular networks and use them as reservoirs to evaluate their cognitive capacity. We further investigate the topological and dynamical underpinnings of their computational properties. Finally, in line with recent work on connectome-based reservoir computing [34, 40, 66, 71, 90, 102, 116, 122, 123, 128, 173, 179, 183], we endow reservoirs with empirical structural connectivity patterns of the human brain to explore the effects of biological hierarchical modularity patterns. To this end, we develop a novel network null model which allows us to disentangle the computational effects of modularity and hierarchical modularity in heterogeneous hierarchical modular networks.

**Figure 1.**
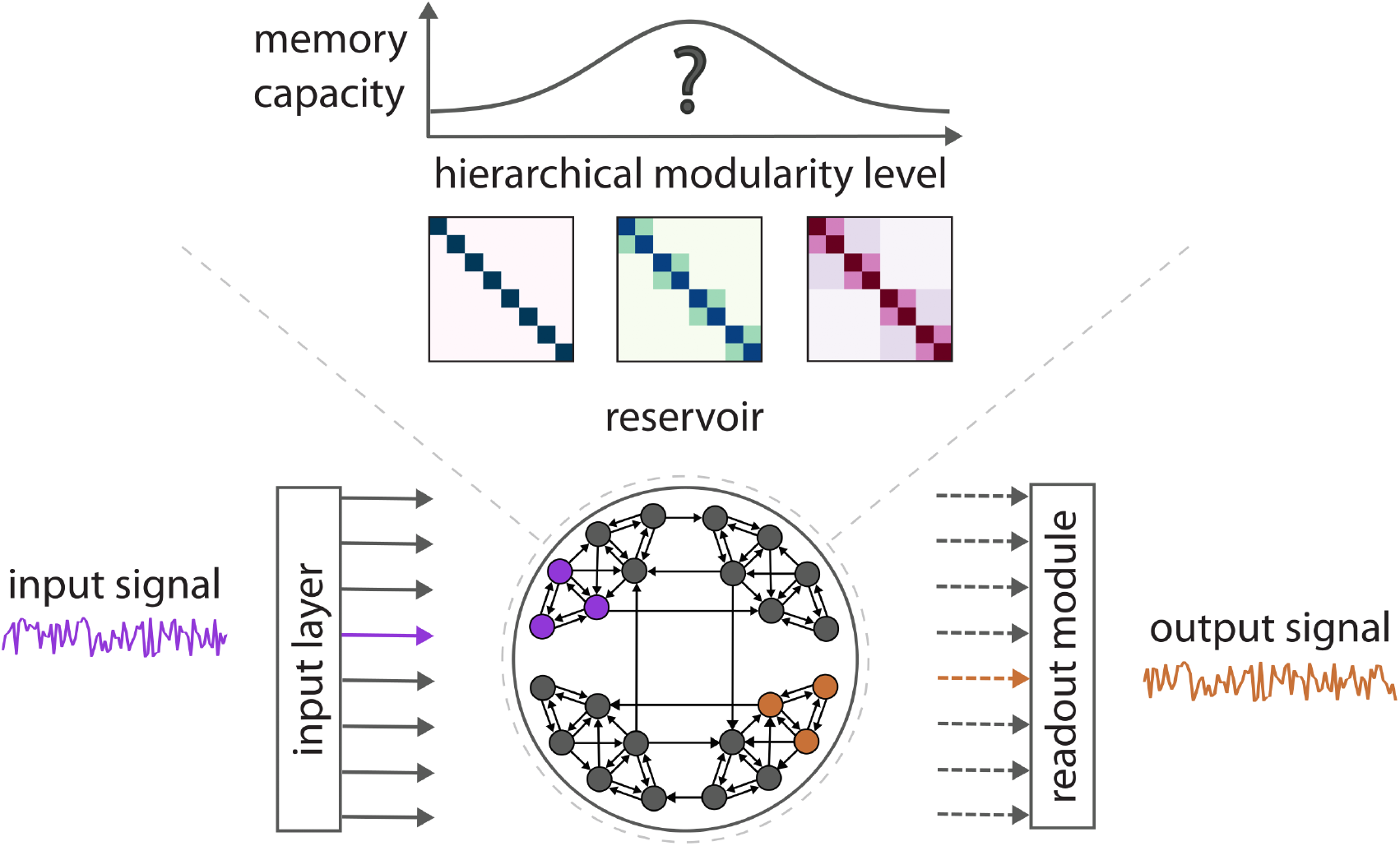
Generating hierarchical modular reservoirs. We study the computational properties of increasing levels of hierarchical modularity using reservoir computing. Top: To evaluate if memory capacity systematically varies with hierarchical modularity, we model a three-level hierarchy bridging strictly modular and hierarchical modular networks. Bottom: We then implement these networks as reservoirs. The reservoir is a fixed recurrent neural network complemented by a readout module. In a learning task, an input signal is introduced via an input module, transformed by the internal dynamics of the reservoir, and recorded from an output module. The readout module is then trained to reproduce a target signal by forming a linear combination of the output signals.

## RESULTS

### Generating hierarchical modular reservoirs

To study the functional effects of hierarchical modularity, we (1) develop a simple method to generate and compare hierarchical modular networks, and (2) implement these networks as reservoirs. We define hierarchical modular networks as networks with a nested blockdiagonal structure. A straightforward method for implementing such networks is stochastic blockmodeling (see *Methods*), which allows us to parametrically tune the number of hierarchical levels, the number of modules at each level, as well as their prominence (modularity). To facilitate systematic comparison between networks with different numbers of hierarchical levels, we develop a method to iteratively remove levels of hierarchy, while preserving basic network features, such as network size, density, degree, and modularity. This ultimately yields a continuum from strictly modular networks to hierarchical modular networks.

Specifically, we model three hierarchical levels of modularity. At the first level, we specify 8 densely connected modules of 50 nodes. We then systematically tune intermodular connectivity to coalesce pairs of lower-order modules into larger, sparser higher-order modules (4 modules of 100 nodes and 2 modules of 200 nodes). For each level, we generate 100 synthetic graphs and randomly assign uniformly distributed weights to the produced edges: ***W*** ∼ *U* (0, 1).

The resulting networks are then implemented as reservoirs to measure their computational capacity (see *Methods*). Briefly, we use a typical reservoir computing architecture in which the reservoir is a recurrent neural network of hyperbolic tangent units, complemented by an input layer and a linear readout module [103]. In a standard learning task, an external time series is introduced into the reservoir through a set of selected input nodes. The input signal propagates across the reservoir and output signals are recorded from a set of selected output nodes. The readout module is then trained using Ridge regression to approximate a target signal by forming a linear combination of the output signals. Importantly, the reservoir remains fixed during training, which enables a clear mapping between the designed network structure and its function. Fig. 1 illustrates the paradigm.

### Hierarchical modularity improves memory capacity

We start by evaluating the reservoir’s ability to preserve representations of past stimuli with the widely used memory capacity task [37, 40, 54, 82, 90, 130, 147, 173, 185] (see *Methods*). In this task, the readout module is trained to reproduce a time-delayed version of a random uniformly distributed input signal. In essence, this amounts to asking the question: To what extent is past input recoverable from the current reservoir states? In contrast to other memory tasks, the memory capacity task isolates memory from other computational capacities like nonlinear processing or pattern recognition. After training the readout, the model’s performance is evaluated on independent data using the ***R***^2^ regression score and summed across a range of time lags to obtain the network’s *memory capacity*. Input nodes correspond to all the nodes in a randomly selected first-level module. All the other nodes in the reservoir are used as output.

Importantly, the computational properties of the reservoir result from an interaction between its structure, which defines the pathways for interaction, and its dynamics, which govern the flow of activity along those pathways. Here, to characterize the relationship between network structure, dynamics, and function, we parametrically tune the reservoir’s global dynamics by scaling its spectral radius *α*. Reservoir dynamics are guaranteed to be stable for *α <* 1 [83, 196]. Conversely, dynamics are expected to be unstable or chaotic for *α >* 1 and are described as critical at *α* ≈1, or at the edge of chaos [54, 173] (see *Methods* and Fig. S1 for more details).

Fig. 2a shows the edge probability matrices used to generate the three levels of the modularity hierarchy (left) and examples of resulting reservoir connectivity matrices (right). In Fig. 2b, we compare the memory capacity of the three network ensembles across different dynamical regimes. For all *α* values considered, we find that performance systematically follows the modularity hierarchy, with higher-order hierarchical modular networks consistently outperforming their lower-order counterparts (*p <* 0.01, common-language effect size (CLES) ≥ 61.15% for all two-tailed, Wilcoxon–Mann– Whitney two-sample rank-sum tests). For completeness, we also show that hierarchical modular networks outperform degree-preserving random networks in Fig. S2a (*p <* 10^−33^, CLES = 100%, two-tailed, Wilcoxon–Mann– Whitney two-sample rank-sum test). For all levels, maximal performance is observed at criticality (*α* = 1) as expected. Altogether, these results demonstrate that hierarchical modularity robustly improves memory capacity over other commonly used architectures across dynamical regimes.

**Figure 2.**
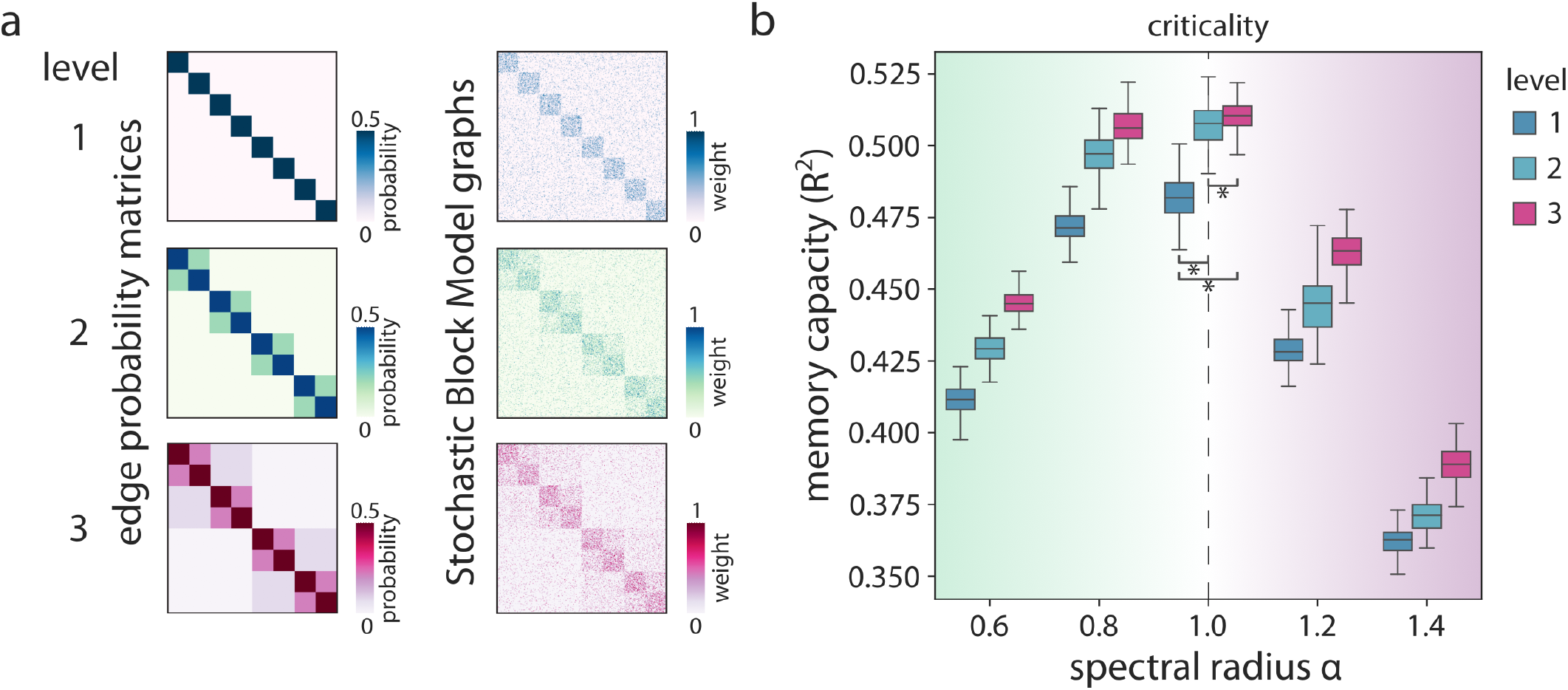
Hierarchical modularity improves memory capacity. (a) Left: Three edge probability matrices, specifying the probability of connecting two nodes based on their community affiliation, were designed to reflect three hierarchical levels of modularity. Right: The three edge probability matrices were used to define three distinct Stochastic Block Models (SBM). Each SBM was used to generate an ensemble of 100 synthetic graphs of which examples are shown. (b) Memory capacity (*R*^2^) as a function of the *α* parameter across hierarchical levels. Asterisks indicate statistical significance of differences between levels according to WilcoxonMann-Whitney tests (*p <* 0.01). Significance is only shown for the highest performing regime.

### Hierarchical modularity generates a pool of timescales

In the previous section, controlling the spectral radius allowed us to relate the reservoir’s structure, dynamics, and function at the global scale. However, how hierarchical modularity shapes local dynamics to improve memory capacity remains unclear. To determine if differences in memory capacity are reflected in node-level dynamics, we compute neural timescales from the nodal activation time series of each reservoir at criticality. The neural timescale is a measure of the speed of decay in the autocorrelation of neural activity. Due to the discrete nature of the reservoir dynamics, timescales are expressed in arbitrary timestep units, but retain the same interpretation as brain-derived timescales, with greater values indicating greater temporal persistence (see *Methods*). In the brain, neural timescales are highly variable [12, 61, 126, 199], both spatially, following a functional specialization gradient from sensory to association cortex [7, 74, 79, 143], and temporally, as they adapt to task requirements [24, 58, 108, 182]. Interestingly, we find no significant difference across hierarchical levels in the reservoir-averaged neural timescales. However, we find that higher-order hierarchical modular reservoirs show more variability in timescales across nodes (Fig. 3, left; *p <* 10^−5^, CLES ≥ 69.98% for all two-tailed, Wilcoxon– Mann–Whitney two-sample rank-sum tests). This reflects a richer temporal expansion of the input signal, which can be visually observed in raw activation time series as a stretching of slow fluctuating signals (Fig. 3, right).

**Figure 3.**
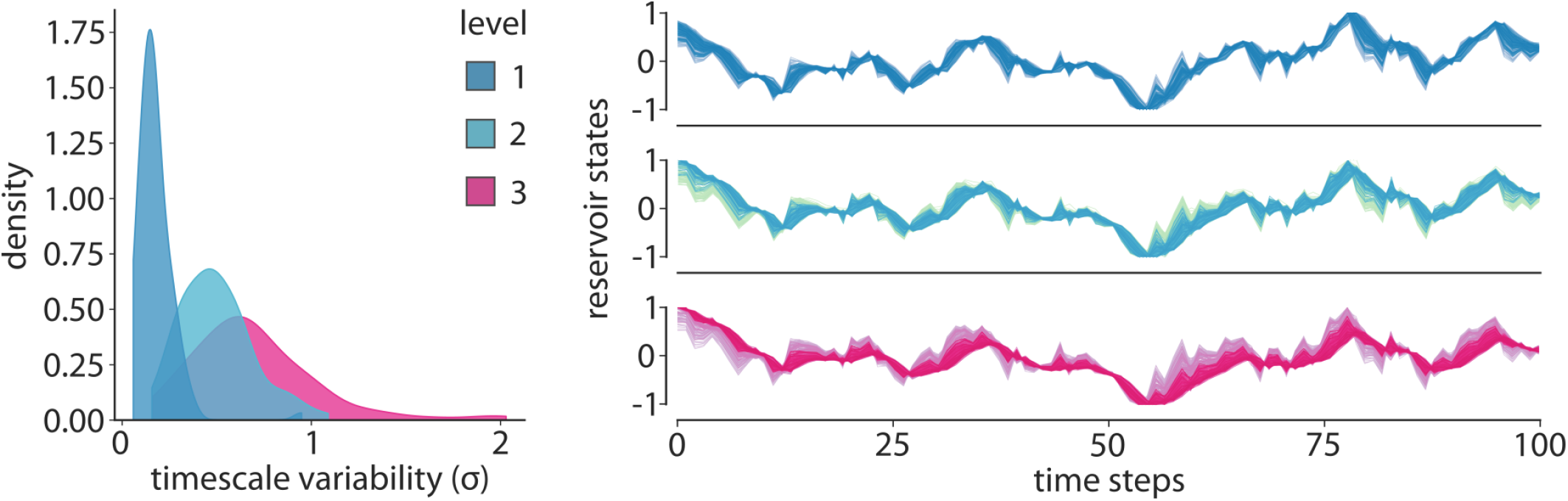
Hierarchical modularity generates a pool of timescales. Left: Distributions of reservoir-wise timescale variability, measured as the standard deviation across neural timescales, for each of the three levels. Right: Output node states at criticality (*α* = 1) for an example reservoir of each of the three levels. The time series were rescaled between −1 and 1 to emphasize differences. Different colours correspond to different nodes.

To uncover the topological underpinnings of these differences in dynamics, we then consider the motif composition of the reservoir. Motifs are local connection patterns that constitute the elementary computational circuits of a network [118, 171]. The motif composition thus tiles a mosaic of recurring functional units, reflecting the network’s computational repertoire. We start by measuring the clustering coefficient, a fundamental measure of a network’s local cohesiveness [52, 192]. Specifically, the network average clustering coefficient measures the prevalence of directed triangles around all nodes. We find that higher-order hierarchical modular networks exhibit a larger clustering coefficient than their lower-order counterparts (Fig. 4a; *p <* 10^−32^, CLES ≥ 98.78% for all two-tailed, Wilcoxon–Mann–Whitney two-sample rank-sum tests), indicative of enhanced local recurrence.

**Figure 4.**
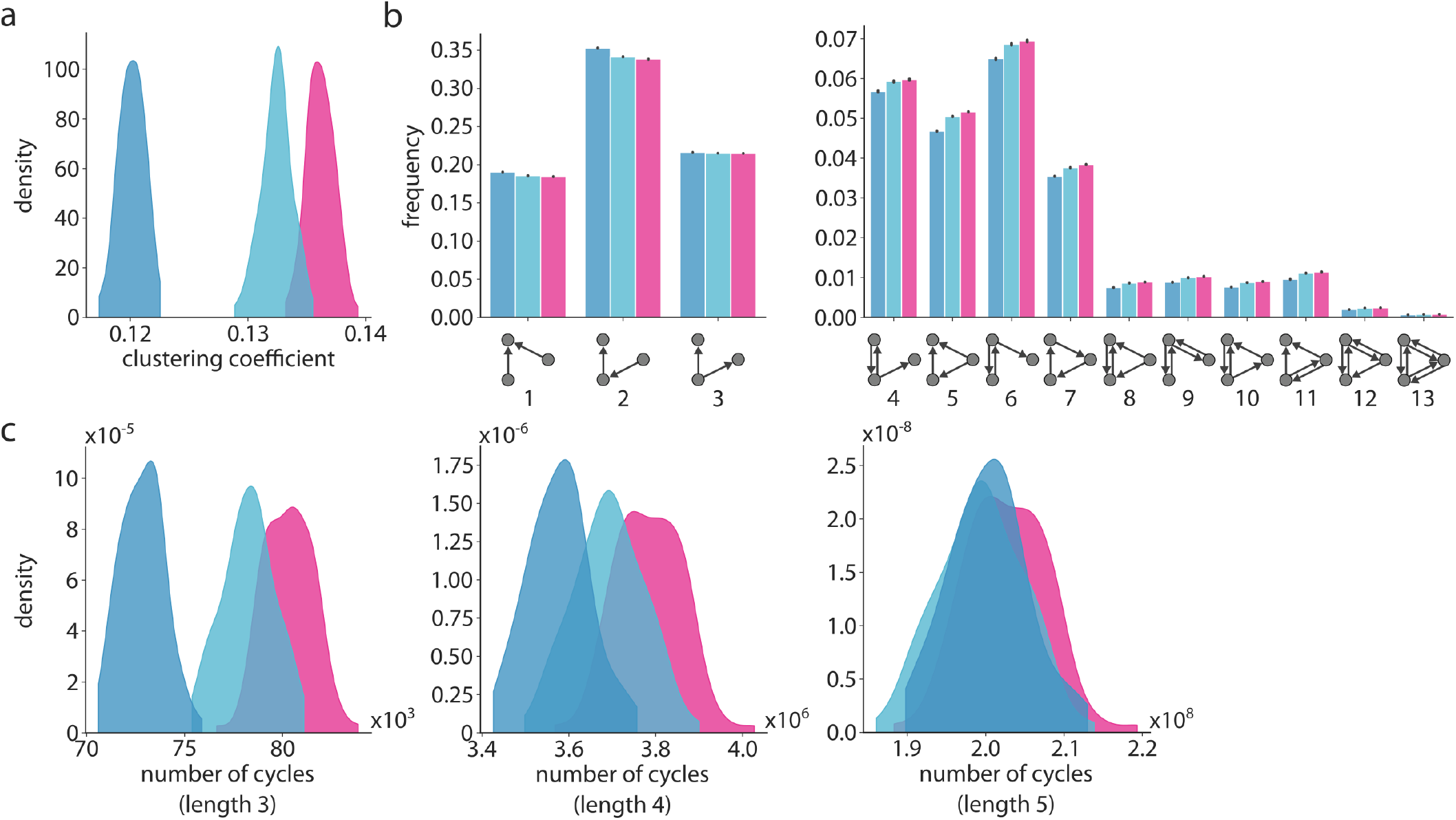
Hierarchical modularity enriches recurrent motifs. (a) Distributions of network average clustering coefficients across hierarchical levels. (b) Motif composition of the reservoirs across hierarchical levels. For all 13 possible three-node directed motifs, bars show the mean frequency across the network ensemble. Error bars correspond to a 95% bootstrapped confidence interval (1000 samples). (c) Distributions of the number of cycles of different length across hierarchical levels.

We then extend our analysis to all possible three-node directed motifs (see *Methods*). For all examined motifs, we find significant differences between all levels of the modularity hierarchy (*p <* 0.05, CLES ≥ 58.64% for all two-tailed, Wilcoxon–Mann–Whitney two-sample ranksum tests), which allow us to partition the motifs into two categories. The three simplest and most prevalent motifs across all networks are all reduced in frequency as levels are added to the hierarchy (Fig. 4b, left). These motifs contain only two edges, with no reciprocal links, and embody basic patterns of convergence (1), sequential processing (2), and divergence (3). Conversely, more complex motifs containing at least three edges are all enriched in higher-order hierarchical modular networks (Fig. 4b, right). These motifs integrate simpler dyadic (two-node) and triadic (three-node) patterns and could therefore subserve more complex computations, notably through reciprocal links and cycles.

In particular, cycles have previously been related to information storage [101] and suggested as a mechanism of time scale separation in modular networks [76, 133]. Therefore, we turn our attention beyond triadic motifs to focus on cycles of varying lengths. Specifically, we consider simple cycles, paths that begin and end at the same node without visiting a node more than once. We find significantly more cycles of length 3, 4, and 5 in higher-order hierarchical modular networks (Fig. 4c; *p <* 0.01, CLES ≥ 62.74% for all two-tailed, Wilcoxon– Mann–Whitney two-sample rank-sum tests, except between levels 1 and 2 for cycle length 5), in line with a richer dynamic repertoire.

Overall, these results point to recurrence and temporal expansion as potential substrates of memory capacity in hierarchical modular networks. To validate this hypothesis, we relate timescale variability and motif composition to memory capacity across all reservoirs (Fig. S3). We consistently find that features related to hierarchical modular reservoirs also relate to enhanced memory capacity.

### Hierarchical modularity supports multitasking

Having evaluated the memory capacity of hierarchical modular networks, we now consider their ability to perform multiple tasks simultaneously. Previous work has suggested that modularity can improve multitasking capacity by providing a balance between inter-modular resource sharing and task interference [102]. Here, we test whether hierarchical modularity can further improve this effect by striking a more robust balance between information integration and segregation [76, 114]. To assess multitasking capacity, we adapt a framework developed by Loeffler et al. [102], which includes a non-linear transformation task, in addition to the memory capacity task. Briefly, the non-linear transformation task consists of regressing a slowly varying sinusoidal input signal to a different waveform — here, a square signal [60, 163]. Beyond evaluating multitasking capacity, this framework also captures the reservoir’s ability to balance two fundamental yet opposing properties: information storage and non-linear information processing [37, 39].

A multitasking experiment proceeds as follows: The set of 4 modules encapsulated in the first of the 2 third-level modules is designated for memory capacity tasks, whereas the other set is assigned non-linear transformation tasks (see Fig. 5a). As for the memory capacity task, input nodes correspond to all the nodes in a randomly selected first-level module. All the other nodes in the third-level module are used as output. Importantly, these task assignments remain consistent across networks spanning all three hierarchical levels. During a task, all input signals are propagated simultaneously across the network and a separate readout module is trained for each task. The multitasking performance is then assessed on independent data as the average ***R***^2^ regression score across all tasks.

**Figure 5.**
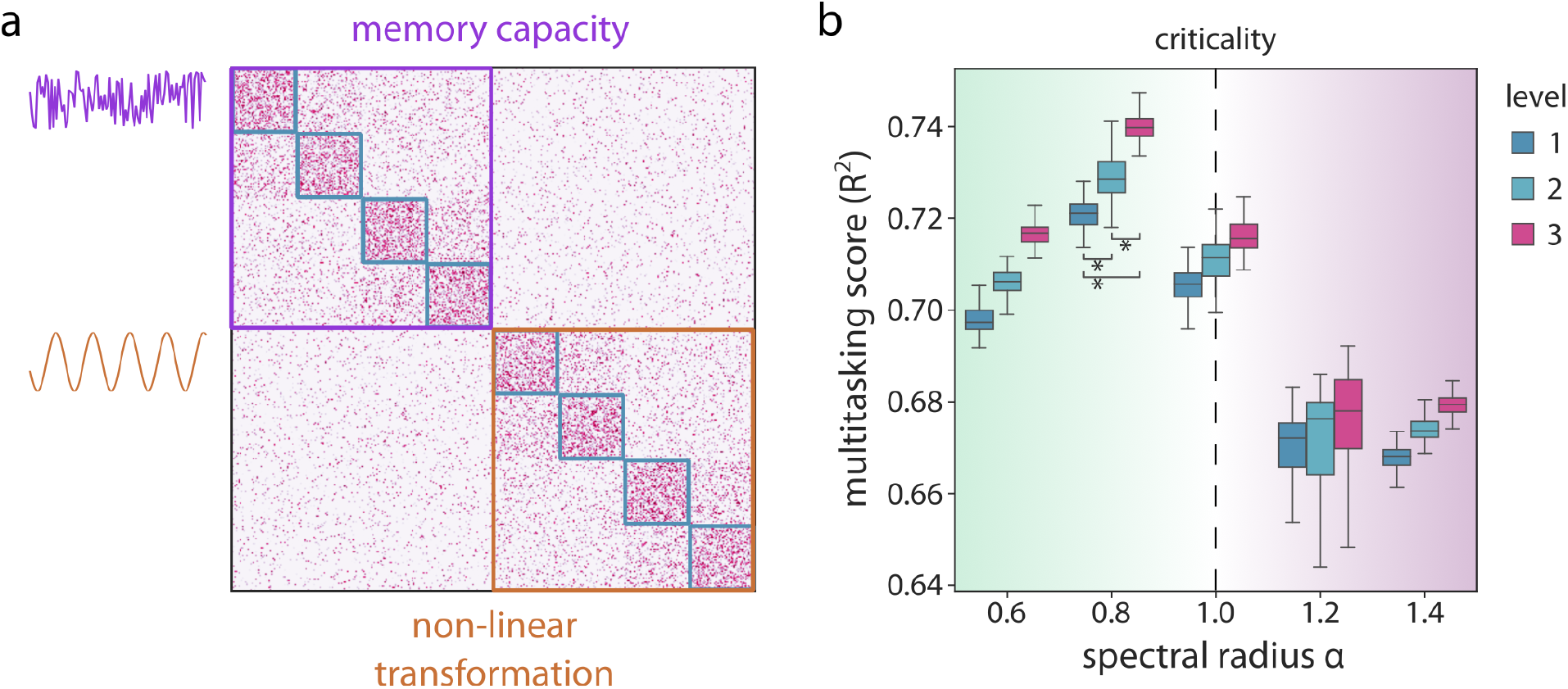
Hierarchical modularity supports multitasking. (a) Schematic of the multitasking framework: the 2 third-level modules are each assigned a different task: the memory capacity task and the non-linear transformation task. Both input signals are propagated simultaneously across the network. (b) Average *R*^2^ regression score across tasks as a function of the *α* parameter for each hierarchical level. Asterisks indicate statistical significance of differences between levels according to Wilcoxon-Mann-Whitney tests (*p <* 10^−25^). Significance is only shown for the highest performing regime.

Fig. 5b shows the multitasking score distributions of the three hierarchical levels of modularity across different dynamical regimes. Again, we find that higherorder hierarchical modular networks consistently outperform their lower-order counterparts (*p <* 10^−25^, CLES ≥ 93.17% for all two-tailed, Wilcoxon–Mann–Whitney two-sample rank-sum tests), including degree-preserving random networks (Fig. S2b; *p <* 10^−33^, CLES = 99.89%, two-tailed, Wilcoxon–Mann–Whitney two-sample rank-sum test). For all levels, maximal scores are observed in a more stable regime (*α* = 0.8) than for the memory capacity task alone. For completeness, we consider multiple other multitasking scenarios which include incorporating more input signals and interleaving task categories instead of assigning them to different third-level modules (Figs. S4, S5; see *Methods* for more details). Across all evaluated scenarios, we find that maximal performance is again observed at criticality (*α* = 1) and that hierarchical modular networks consistently outperform strictly modular ones. Note however that the third level of hier-archy rarely leads to additional improvements over the second level. Altogether, these results demonstrate that hierarchical modularity robustly supports multitasking in various experimental scenarios. Importantly, since performance rapidly saturates with additional hierarchical levels, our findings also suggest that a deep fractal organization might not be necessary to achieve performance improvements.

### Hierarchical modularity in the human connectome

Until now, we have considered synthetic blockmodel graphs that are ostensibly neuromorphic, in so far as they reflect the multi-scale nested organization of brain modules [8, 14]. However, these networks lack the heterogeneity of empirical brain connectomes, that is, realworld modules vary in size and in the number of submodules that they contain. To account for these differences, we endow reservoirs with empirical structural connectivity patterns of the human brain. Specifically, we build a group-representative connectome from 327 individual structural networks derived from diffusion-weighted magnetic resonance imaging (MRI) tractography (source: Human Connectome Project - HCP [184]; see *Methods* for detailed procedures).

To isolate the effect of hierarchical modularity on the performance of connectome-informed reservoirs, we develop a novel hierarchical modular network null model. Our model draws on an existing modular random graph model [153] and the intuitive notion that modularity can be operationalized as the relative density of intramodular connections [59, 127]. For a strictly modular network null model, we can distinguish two categories of edges: within- and between-module edges. We can then rewire them separately while constraining the edge swap so that the rewired edges retain their original category. This allows us to randomize the network while preserving its modularity. Generalizing to hierarchical modularity, we can infer that *L* nested partitions will result in *L* + 1 edge categories in order for each module of each partition level to maintain intra-modular connectivity (see *Methods* for more details on how these categories are identified). Similar to the procedure used in the modular null model, we then rewire edges of each category separately, using the classic switching method [111]. The resulting null network’s connectivity patterns are randomized while maintaining its size, density, degree sequence, and hierarchical modularity.

To assess the computational properties of realistic hierarchical modularity, we start by applying Louvain modularity maximization to the empirical connectome. This method naturally yields a hierarchical partition: each node is assigned to a single module at each level and lower-level modules can be nested within higher-level modules. Since the algorithm is non-deterministic, we re-run it multiple times and select a representative solution (see *Methods* for more details). The retained solution has 2 hierarchical levels with 15 modules at the first level and 6 modules at the second level (Fig. 6a). Only one of the first-level modules is not nested into a second-level module. Next, using the derived partition, we apply both the modularity-preserving and the hierarchical modularity-preserving null models to the connectome to generate two null network ensembles of 100 networks each. Fig. 6b shows the within- and between-module edge categories which constrain rewiring in each model (left), as well as examples of resulting null networks (right), highlighting their preserved modular and hierarchical modular architectures, respectively.

**Figure 6.**
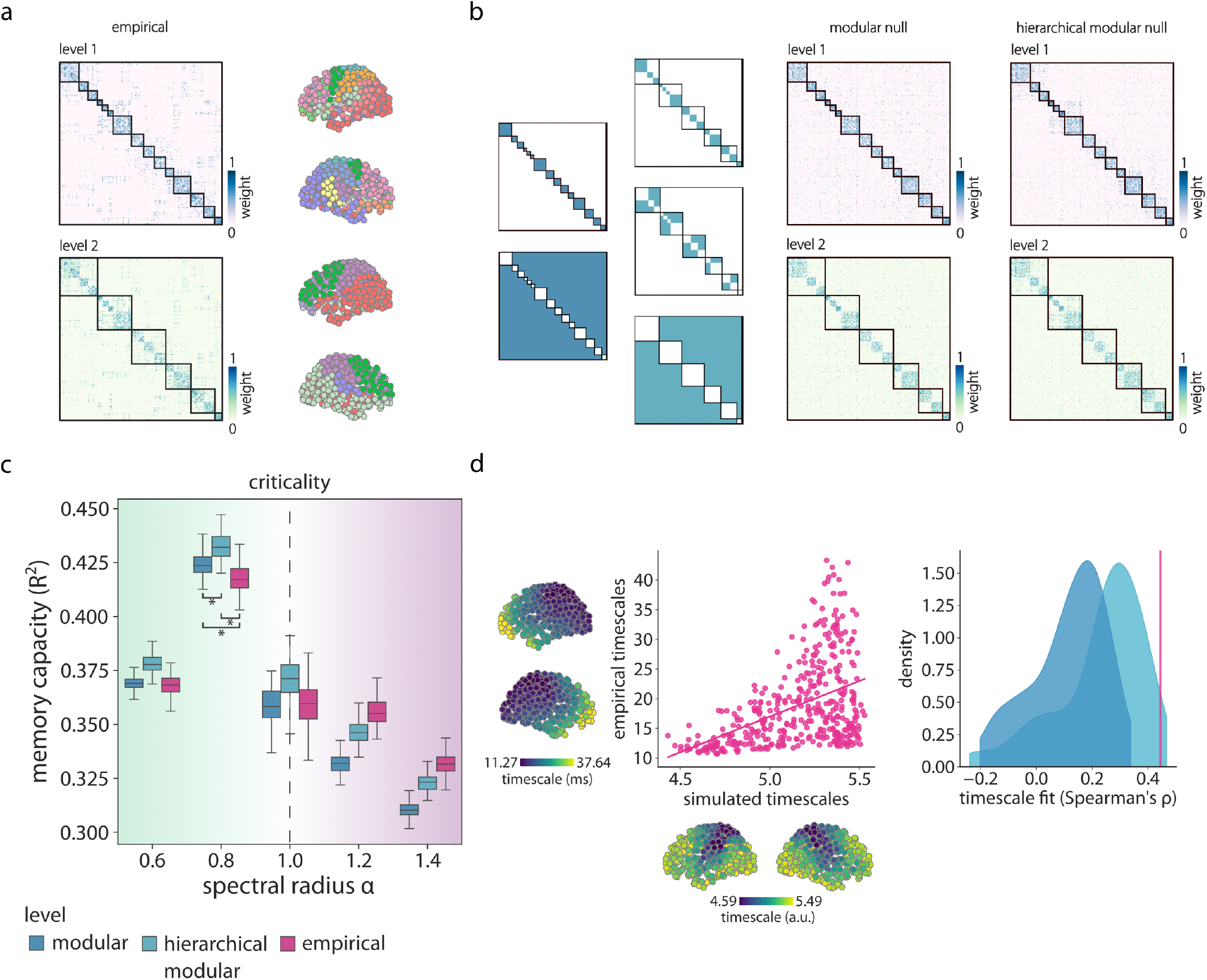
Hierarchical modularity in the human connectome. (a) Left: Empirical human brain structural connectivity matrices. Borders delineate the communities of a two-level hierarchical partition identified using Louvain modularity maximization. The second-level partition (bottom) encapsulates the first-level partition (top). Right: Brain mapping of the modules at each level. Each colour represents a different module. (b) Left: Edge categories constraining the modularity-preserving (in blue) and hierarchical modularity-preserving (in teal) rewiring procedures. Right: Example modularity- and hierarchical modularity-preserving null networks with the empirical module borders overlayed to showcase their preserved (hierarchical) modular structures. (c) Memory capacity (*R*^2^) as a function of the *α* parameter across hierarchical levels. Asterisks indicate statistical significance of differences between levels according to Wilcoxon-Mann-Whitney tests (*p <* 10^−9^). Significance is only shown for the highest performing regime. (d) Left: Relationship between empirical and simulated timescales. Each point represents a brain region. A linear regression line is shown for visualization purposes. Right: Null distributions of timescale fits measured as the Spearman correlation coefficient between empirical and simulated timescales. Null fits obtained using the (hierarchical) modularity-preserving null networks are shown in blue (teal). The fit obtained from the empirical connectome is shown in magenta.

Next, we instantiate reservoirs using the null connectivity matrices, as well as the empirical connectome, and use them to perform the memory capacity task. As for the synthetic networks, first-level modules are used as inputs and outputs. Performance is averaged over all possible pairwise combinations of inputs and outputs. We use 100 different input signals for the empirical connectome to create a performance distribution comparable to those of the null ensembles. Fig. 6c shows the memory capacity distributions of all three ensembles as a function of spectral radius (*α*). Across all dynamical regimes, we find that the hierarchical modular null networks significantly outperform their strictly modular counterparts (*p <* 10^−12^, CLES ≥ 79.5% for all twotailed, Wilcoxon–Mann–Whitney two-sample rank-sum tests), in line with the results observed in the homogeneous hierarchical modularity model. Interestingly, maximum performance is observed in a more stable regime (*α* = 0.8), consistent with previous reports of memory capacity in connectome-informed reservoirs [173]. Moreover, in the stable regime (*α <* 1), the hierarchical modular null ensemble consistently outperforms the empirical connectome (*p <* 10^−24^, CLES ≥ 92.82% for both two-tailed, Wilcoxon–Mann–Whitney two-sample ranksum tests), suggesting that in this regime, additional topological features of the human brain are detrimental to maintaining signal representations.

To further characterize this variability in topological features beyond modularity, in Fig. S6, for each module used as input (output), we average memory capacity over all possible outputs (inputs). This results in a measure of how effective a module is as an input (output). We then relate module-wise performance to various module-average weighted nodal features, namely strength (sum of edge weights incident to each node), clustering (average intensity of triangles around a node [129]), and node-average communicability (communication measure integrating all possible walks on a network [35, 50]) using Spearman correlation. We find that both input and output performance is strongly negatively related to clustering (input: *ρ* ≈™0.6, *p* ≈0.02, output: *ρ* ≈™0.62, *p* ≈0.01). Output performance is also strongly positively related to strength (*ρ* ≈0.56, *p* ≈0.03) and communicability (*ρ* ≈0.68, *p* ≈0.006). Collectively, the results indicate that segregated input and output modules reduce memory capacity. In particular, highly-performing output modules are also highly integrated in the network, as indicated by high strength and communicability.

Finally, to evaluate the realism of connectome-informed reservoir dynamics, we contextualize them against empirical brain timescales. Specifically, we average output node timescales in the memory capacity task across all 100 input signals and 15 input modules. We then compare the resulting connectome-informed synthetic timescale map with an empirical brain map of group-averaged resting-state intrinsic timescales derived from magnetoencephalography (MEG) data acquired in a subset of 33 participants from the HCP (see *Methods* for additional information) [157]. We find a significant positive relationship between the synthetic and the empirical timescales using Spearman correlation (Fig. 6d, left, *ρ ≈* 0.44, *p <* 10^−20^). Importantly, we also contrast the empirical correlation coefficient with null correlation coefficients obtained using the synthetic timescales derived from modularity- and hierarchical modularity-preserving null reservoirs (Fig. 6d, right). We define a two-sided p-value (*p*_*rewired*_) as the proportion of more extreme null coefficients. We find that the empirical relationship is significantly greater than the null relationships obtained from modularity-preserving reservoirs (*p*_*rewired*_ *<* 0.01) but not hierarchical modularity-preserving reservoirs (*p*_*rewired*_ ≈ 0.13). Together, these results show that hierarchical modularity could contribute to shaping empirical gradients of intrinsic neural timescales in the brain.

### Sensitivity analyses

By replicating our findings using empirical brain connectivity patterns and (hierarchical) modularity-preserving network null models, we have shown that our results generalize to heterogeneous modularity patterns. Here, we further assess the sensitivity of the results to several other modeling choices, namely the edge probability scaling factor *r* (see *Methods* for more details), the size of the networks, their symmetry, the sign of the connection weights, and the presence of self-loops (connection from a node to itself). In the main analysis, we used asymmetric (directed) networks of *N* = 400 nodes, with positive weights and no self-loops. Inter-modular connectivity was halved (*r* = 0.5) at each hierarchical level. In Fig. S7, we additionally test *N* -values of 200 and 600 nodes, and *r*-values of 0.25 and 0.75. We find that hierarchical modular networks systematically outperform their strictly modular counterparts, with peak performance near criticality (*α* = {0.8, 1.0}), independently of the network sizes and scaling factors considered. In Fig. S8, we show that this result also extends to symmetric (undirected) networks and networks containing self-loops. However, no significant difference was found between levels of the modularity hierarchy when including negative weights (***W*** ∼*U* (− 1, 1)). Furthermore, asymmetric networks including negative weights can achieve near perfect performance at high *α* values. Nevertheless, we find that introducing negative weights only in within-module connections restores the effect of hierarchical modularity, despite a drop in performance compared to allowing negative weights throughout the network. Altogether, these results confirm that the computational advantages of hierarchical modularity are robust to numerous modeling settings and suggest it could constitute a versatile design principle for artificial neural networks.

## DISCUSSION

In the present report, we develop a simple blockmodeling method to generate and compare multi-level hierarchical modular networks. We implement these networks as reservoirs to evaluate their computational properties. We find that hierarchical modular networks exhibit greater memory capacity than strictly modular networks. This performance increase is associated with more diverse neural timescales in hierarchical modular reservoirs and a higher prevalence of reciprocal and cyclic motifs. Furthermore, we show that the computational advantage of hierarchical modularity extends to a multitasking setting, as well as heterogeneous empirical structural connectivity patterns. Finally, using a novel hierarchical modularity-preserving network null model, we find that hierarchical modularity might play an important role in shaping the diverse portrait of intrinsic neural timescales we observe in the brain.

These results extend previous work on the benefits of modular and hierarchical architectures in reservoir computing [85, 102, 113, 147, 195]. In particular, Rodriguez et al. [147] found that modular networks of threshold neurons showed improved memory capacity in contrast to random networks. They suggest that this effect is obtained by balancing local modular cohesion and global communication to achieve maximal network activity. However, they found modularity to decrease memory performance when using a hyperbolic tangent activation function. Here, we show that hierarchical modular networks can outperform both strictly modular and random networks at the memory capacity task in classic reservoirs of hyperbolic tangent neurons. This effect might be due to hierarchical modularity providing a more robust balance of segregative and integrative graph features through a principled scaling of connectivity from the local to the global level [76].

Such an architecture can also prove useful for multitasking, as we show under various experimental scenarios combining memory capacity and non-linear transformation tasks. These results extend previous work from Loeffler et al. [102], who showed that modularity improved multitasking performance in neuro-memristive nanowire networks. By taking advantage of modules for input and output placement, hierarchical modularity might maximize resource allocation via integration while limiting task interference via segregation [102]. Alternatively, a certain level of integration might also benefit generalization via regularization by noise from different tasks [17, 23]. More generally, recent work has emphasized a fundamental tradeoff between network architectures supporting parallel processing of independent tasks and interactive processing of tasks with shared representations [139]. Future research could explore the potential of hierarchical modular networks to balance these competing demands — leveraging shared representations to facilitate generalization [23], while limiting interference to support multitasking in tasks with overlapping structure. More broadly, these results open the door to tailoring network architectures to the structure of specific task domains. In particular, hierarchical modularity might prove useful in tackling more complex compositional task structures. In line with this hypothesis, it was previously shown that hierarchical modular architectures could be the expression of an evolutionary pressure to adapt to a rapidly changing environment by dividing complex changing goals into reusable basic subtasks [89].

The improved performance of hierarchical modular networks could also be understood through the lens of their local topology. Notably, additional levels of modules increase the prevalence of cycles of different lengths. These connection patterns could act as storage circuits [101], maintaining the input signal over diverse time horizons [62]. They can also constitute a tuning mechanism for a reservoir’s frequency response [1]. More generally, by analyzing all possible three-node motifs, we find that hierarchical modular networks are enriched in complex reciprocal and cyclic motifs. These results extend previous work from Sporns [168] et al. characterizing the motif composition of fractal connectivity patterns. Of note, the enrichment of motif 9 (see Fig. 4b) in hierarchical modular networks echoes consistent results across numerous large-scale mammalian brain networks [100, 171]. This motif consists in a chain of reciprocally connected nodes that do not form a loop. It denotes the potential for strong signaling between certain node pairs and the absence of direct connections between others. For this reason, it has been posited as another potential mechanism subserving the balance between information integration and segregation [100, 171]. Altogether, these results reiterate the importance of considering higher-order interactions to understand the behaviour of real-world systems [9, 19].

From a dynamical perspective, modular networks have previously been shown to give rise to time-scale separation, allowing to distinguish fast intra-modular and slow inter-modular processes [133]. These results have been generalized to hierarchical modularity in oscillator networks, showing the emergence of as many timescales as there are hierarchical levels [6, 165]. Here, we study neural timescales in hierarchical modular networks as the decay time of a node’s activity autocorrelation, similar to how it is studied in the brain [61, 126]. We find an increase in the diversity of neural timescales in higherorder hierarchical modular networks, which positively correlates with their improvement in memory capacity. This result adds to accumulating computational evidence that heterogeneous and hierarchically organized timescales enhance the processing of complex temporal information [30, 64, 70, 93, 109, 124, 137, 142, 176, 197].

Beyond functional implications, our results in connectome-informed reservoirs complement a growing body of computational modeling work exploring the emergence of a timescale hierarchy in the brain. While biophysical approaches have mostly focused on time constants associated with cellular and synaptic processes, network-based accounts offer important complementary explanations [47, 64, 199]. Notably, clustered connectivity patterns have been associated with the emergence of slow timescales [80, 98]. Some models, such as the one by Chaudhuri et al. [27], also offer integrative accounts, combining neurobiologically grounded dynamics and connectivity. Using a threshold-linear recurrent network constrained by empirical macaque inter-regional connectivity, the authors show that both long-range projections and a cytoarchitectural gradient of excitation shape the emergence of a cortical timescale hierarchy. Here, we go beyond the qualitative matching of timescales to gradients of brain organization and directly relate simulated timescales to empirical neural timescales derived from MEG. We find that while our model relies solely on structural connectivity and generic nonlinear units, it can still produce a rich landscape of timescales reminiscent of empirical observations. Furthermore, using a novel hierarchical modularity-preserving network null model, we show that hierarchical modularity may constitute a core architectural principle underlying intrinsic timescales, offering a parsimonious account that complements biologically detailed models. Overall, these results reinforce the idea that network topology is not just a passive substrate, but an active determinant of neural dynamics, shaping computational capacity through structured heterogeneity.

From a modeling perspective, this work introduces two main novelties. First, we develop an intuitive stochastic blockmodeling method to generate and compare hierarchical modular networks. Numerous models of hierarchical modularity have already been developed [6, 21, 85, 87, 88, 121, 132, 144, 146, 152, 154, 168, 190, 198]. Notably, Moretti et al. [121] distinguish a bottom-up approach — local modules are recursively connected with level-dependent density [121, 132, 168] — and a top-down approach — starting from a homogeneous random network, modules are recursively split and intermodular connections are rewired into their source module with a given probability [190]. In contrast, our model directly defines the whole nested block modular architecture via an edge probability matrix, with a size determined by the number of modules at the first level of the hierarchy. Previous modeling approaches mainly focused on an edge probability scaling parameter, bridging disconnected modular networks and homogeneous random networks [168] or small-world networks [121]. Here, we introduce a simple and principled approach that begins with hierarchical modular networks of arbitrary depth and incrementally removes hierarchical levels, creating a continuum between strictly modular and hierarchical modular architectures while preserving network size, density, degree, and modularity. Second, we introduce a novel, broadly applicable network null model that randomizes network topology while preserving network size, density, degree, and hierarchical modular structure. This model can be applied to any complex hierarchical modular network to disentangle topological or dynamical effects resulting from higher-order topological constraints from those passively endowed by hierarchical modularity. In network neuroscience, this work aligns with a growing effort to develop increasingly realistic network null models that retain meaningful structural properties beyond basic constraints such as degree or density [117, 149, 150]. An important benchmark model in network neuroscience combines regional connectivity (degree) and modules [150, 169]. Future work could explore if the addition of a hierarchical modular architecture holds significantly more explanatory power in recapitulating brain network organization.

The present findings should be interpreted with respect to some methodological limitations. First, to isolate the effects of hierarchical modularity on memory capacity, we employ a classic reservoir model equipped with simple, homogeneous dynamics. Future studies could examine the effect of heterogeneous dynamics and introduce incremental layers of biological details, including microarchitectural excitation gradients [27, 43], cell type-specific dynamics [180], or geometry-informed conduction delays [164]. Second, we find that the effect of hierarchical modularity on memory capacity only generalizes to signed networks when negative weights are introduced in first-level modules. This indicates that random placement of negative weights might effectively blur a binary hierarchical modular organization. Given the importance of negative connections for representation capacity [188], future work should explore how to most efficiently integrate them in hierarchical modular architectures. Interestingly, placing negative weights only in low-level modules reflects the prevalent modeling assumption that the brain is organized in local populations of interacting excitatory and inhibitory neurons, coupled exclusively through excitatory long-range connections [193, 194]. More broadly, these results pave the way for more complex multi-region “network of networks” RNN models [138]. Finally, the structural connectome used in this study was reconstructed using diffusion-weighted MRI, which is prone to false positives and false negatives [41, 106, 178]. Although we mitigated this by constructing a group-representative connectome, future work could benefit from using data derived from more accurate invasive techniques such as tract tracing to validate and extend our findings.

In summary, this work introduces a principled framework for constructing and comparing hierarchical modular networks, demonstrating that hierarchical modularity can enhance memory capacity, multitasking, and temporal processing in neuromorphic reservoir systems. Extending these findings to connectome-informed reservoirs, we show that hierarchical modularity contributes to the emergence of brain-like neural timescales. Together, these results advance our understanding of structure-function relationships in neural systems and point to hierarchical modularity as a powerful inductive bias for the design of neuromorphic computing architectures. Future work could build on this foundation by exploring more biologically realistic dynamics, and validating these principles in real neural circuits. More generally, this study contributes to ongoing efforts at the intersection of neuroscience, artificial intelligence, and cognitive science, demonstrating how network science may serve as a shared language for advancing crossdisciplinary knowledge.

## METHODS

Code and data used to perform the analyses can be found at https://github.com/netneurolab/milisav_hierarchical_modularity.

### Hierarchical modular network model

Here, we propose a model for constructing homogeneous hierarchical modular networks using stochastic blockmodeling. In contrast to other models of hierarchical modularity, our implementation allows for systematic comparison between networks with different numbers of hierarchical levels, while preserving basic network features, such as network size, density, degree, and modularity.

We start by defining hierarchical modular networks in line with the definition of “networks with a nested hierarchical organization” proposed by Sales-Pardo et al. [154]. First, we define modules as communities of nodes that are more densely connected to each other than to other nodes in the network. Each module’s internal connectivity can, in turn, be divided into smaller submodules at increasingly lower, more fine-grained hierarchical levels. Importantly, submodules are entirely determined by the connections between nodes of the higher-order module, i.e., they are encapsulated [76, 154]. Note that this definition does not accept the existence of soft boundaries or overlap between modules. Furthermore, in line with Sales-Pardo et al. [154], we restrict our model to homogeneous hierarchical modular networks, that is, at each hierarchical level, all modules have the same size and can be divided into the same number of lower-order modules.

From this definition, we can infer that the resulting networks will have a nested block-diagonal structure [146, 154]. They can therefore be implemented using a stochastic block model [21, 146], where edges are added independently for each pair of nodes, with a probability that only depends on the blocks to which the nodes belong [78]. The model is implemented as follows: Consider *M*_*l*_, the number of modules at level *l*, each of size *n*_*l*_ nodes. Next, consider *g*_*l*_ = *M*_*l*_ ^−^_1_*/M*_*l*_ = *n*_*l*_*/n*_*l*_ ^−^ _1_ for *l >* 1, the number of modules at level *l* − 1 encapsulated in each module at level *l*, with *L*, the total number of hierarchical levels. We start by creating an empty edge probability matrix of size *M*_1_ ×*M*_1_. This matrix specifies the density of edges within (diagonal) and between (off-diagonal) blocks (modules) of the first hierarchical level. First, we fill the diagonal with *p*_1_, the probability that two nodes are connected if they belong to the same module at level 1. Second, for each consecutive set of *g*_2_ blocks, we assign *p*_2_ (the probability that two nodes are connected if they are part of different modules at level 1, but the same module at level 2) to the off-diagonal blocks that complete a square of size *g*_2_ blocks. We repeat this step for each hierarchical level, with 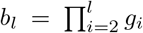 the number of diagonal blocks encapsulated in each level-*l* module. The final level consists of the whole network. Importantly, given that modules must have greater internal than external density, *p*1 *> p*2 *>* · · · *> p*_*L*_. Here, we define the edge probability scaling as *p*_*l*_ = *r*^*l*−1^*p*_1_, with 0 *< r <* 1. Together, the resulting edge probability matrix and *n*_1_ fully determine the model.

From this, we can derive the expected degree *E*[*d*] of a node as the sum of contributions from all levels. At level 1, each node can connect to *n*_1_ − 1 other nodes in its module (excluding self-loops) with intra-modular probability *p*_1_. Therefore, the expected contribution at level 1 is:

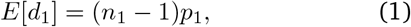

At higher levels *l* ≥ 2, each node can connect to nodes in the other *g*_*l*_ −1 submodules within its module. The number of available connections is therefore (*g*_*l*_ −1)*n*_*l*_ −_1_. With probability *p*_*l*_ = *r*^*l*−1^*p*_1_, the expected degree contribution is:

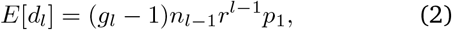

Summing over all levels:

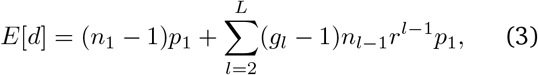

Using the recursive formula 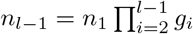, we obtain:

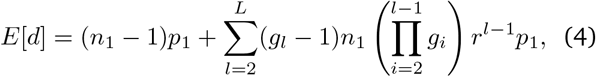

Here, we construct hierarchical modular networks with *M*_1_ = 8 first-level modules of *n*_1_ = 50 nodes, with intra-modular edge probability *p*_1_ = 0.5. These modules are iteratively paired (*g*_*l*_ = 2) into higher-order modules, with inter-modular connectivity *p*_*l*_ halved (*r* = 0.5) at each hierarchical level, for a total of *L* = 4 hierarchical levels. This yields the following edge probability matrix:

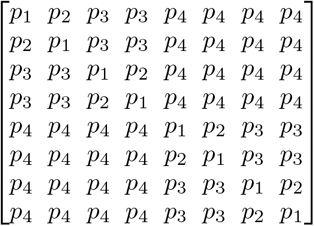

Next, we develop a method for comparing networks with different numbers of hierarchical levels, while preserving basic network features, such as network size, density, degree, and modularity on average. Starting from a network with *L* hierarchical levels, levels can be iteratively removed, starting from the highest level *L*. This is done be replacing, in the edge probability matrix, the values *p*_*L*_ and *p*_*L*−1_ by:

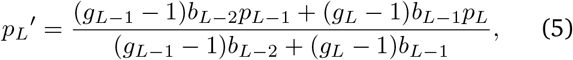

with *p*_*L*_^*′*^, the new inter-modular connectivity at the highest level *L*^*′*^ = *L* − 1, resulting from the average of *p*_*L*_ and *p*_*L*_^−^ _1_, weighted by their prevalence in each row (or column) of the edge probability matrix.

This procedure maintains the expected degree, and by extension, number of edges of the resulting network fixed. Moreover, since *M*_1_ and *n*_1_ are maintained, network size and density are also preserved. Finally, since the edge probabilities within modules at the preserved hierarchical levels are maintained, and the row (or column) sums of the edge probability matrix are maintained, modularity, as measured using Newman’s Q [127], is also preserved for the remaining hierarchical levels.

Here, we run this procedure twice, bridging strictly modular and hierarchical modular networks across three levels. We implement all stochastic block models in the hierarchy using the openly available NetworkX package (https://networkx.org/documentation/stable/index.html) [72]. They all share an average degree of 62 and a density of 15.5%.

### Reservoir computing

To relate hierarchical modular network structure to function, we use reservoir computing, a dynamical systems-oriented machine learning framework taking its roots in computational neuroscience [45, 84, 103, 105]. In a classic reservoir computing architecture, the reservoir is a non-linear recurrent neural network, complemented by an input layer and a linear readout module. In a standard learning task, an external time series is introduced into the reservoir through a set of selected input nodes. The input signal propagates across the reservoir and output signals are recorded from a set of selected output nodes. The readout module is then trained to approximate a target signal by means of a linear combination of these output signals. Importantly, the reservoir remains unchanged during training, which makes this model very efficient and facilitates the mapping of the reservoir’s architectural features to its performance. Fig. 1 illustrates the paradigm.

Reservoir computing relies on two fundamental computational properties: *fading memory* — reservoir states depend only on a finite history of past inputs — and *pairwise separation* — distinct input histories lead to distinct network states [37, 96, 105]. The reservoir’s recurrent connections passively endow it with memory, whereas the complex non-linear interactions that govern its dynamics perform a high-dimensional temporal expansion of the input signal, which has the potential to transform non-linearly separable patterns into linearly separable representations [37].

#### Reservoir dynamics

The reservoir is a recurrent neural network of hyperbolic tangent units. The reservoir states obey the following discrete-time update equation:

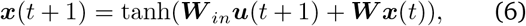

where ***x***(*t*) is the vector of nodal reservoir activation states at time *t*, ***u***(*t*) is the input signal at time *t*, ***W*** _*in*_ is the binary input matrix mapping the input signal to the input nodes, and ***W*** is the reservoir weight matrix. For the stochastic block model graphs, we assign random uniformly distributed weights to the produced edges: ***W*** ∼ *U* (0, 1). For the empirical connectome, weights correspond to the connectivity measures derived from diffusion MRI data.

#### Stability

We parametrically tune the global dynamics of the reservoir by fixing its spectral radius, i.e., the modulus of its leading eigenvalue, to desired values. Specifically, the weight matrix was divided by its spectral radius, effectively fixing it at 1. The resulting matrix was then multiplied by a range of *α* values (*α* = {0.6, 0.8, 1.0, 1.2, 1.4}), fixing its spectral radius at *α*; ***W*** in equation 6 can therefore be expressed as:

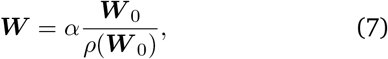

where ***W*** _0_ is the original weight matrix and *ρ*(***W*** _0_) is its spectral radius. Reservoir dynamics are guaranteed to be stable for *α <* 1^1^ [83, 196] and therefore satisfy the *echo state property* for all inputs — a property similar to *fading memory* in which state trajectories converge for the same input independently of initial conditions. Conversely, dynamics are unstable or chaotic for *α >* 1 and are described as critical at *α* ≈1, or at the edge of chaos [54, 173]. Note however that dynamics are not always chaotic for *α >* 1. Notably, recurrent neural networks can be stabilized by strong inputs [49, 103, 120], which can lead to unit saturation [37, 39, 130, 196]. In our experiments, strictly positive weights also seem to lead to unit saturation (see Fig. S1 for Lyapunov exponent estimations using the method presented in [187]).

#### Readout module

The readout module approximates a target signal ***y***(*t*) by means of a linear combination of selected output signals ***x***_*out*_(*t*):

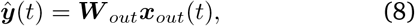

where ***W*** _*out*_ is the readout weight matrix and ***ŷ*** is the final output which approximates ***y***(*t*). The weights are trained in a supervised setting using Ridge regression as implemented in sklearn (https://scikit-learn.org/stable/) [136].

### Memory capacity

To evaluate reservoir memory, we chose the widely used memory capacity task, which measures the reservoir’s ability to preserve the representations of past stimuli in continuing iterations of its internal state computation [37, 40, 54, 82, 90, 130, 147, 173, 185]. In this task, the readout module is trained to reproduce a time-delayed version of a random uniformly distributed input signal ***u***(*t*) ∼*U* (−1, 1). Specifically, ***y***(*t*) = ***u***(*t* −*τ*), where *τ* is a given time lag. Here, we consider 16 time lags, monotonically increased in one time point steps in the range [1, 16]. Note that there are as many linear units as there are time lags in the readout module and they are all trained independently.

We separately generated a training and a testing input signal of 2100 time points each. Different signals were used for each network evaluated. Reservoir states were simulated separately for each timeseries. The first 100 time points were discarded from the resulting reservoir state trajectories to account for initial transients.

The training reservoir states of the selected output nodes were then used to train the readout module to reproduce the input signal at different time lags. Finally, the testing reservoir states were used to test the performance. For every *τ*, performance was measured using the ***R***^2^ coefficient of determination regression score. Regression scores were then summed across all time delays to obtain the memory capacity (MC):

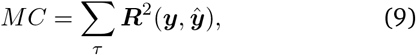

For the synthetic hierarchical modular networks, input nodes correspond to all the nodes in a randomly selected first-level module. Output nodes correspond to the nodes of all the other 7 modules. A different readout module is trained for each one and final performance is averaged across all readout modules. The 8 first-level modules are used as input and output across all levels of the hierarchy. For the empirical connectome, performance is averaged across all possible combinations of input and output modules to account for module heterogeneity.

### Multitasking

To evaluate reservoir multitasking capacity, we adapt a multitasking framework developed by Loeffler et al. [102] for use in neuro-memristive nanowire networks. The multitasking setup involves the memory capacity task, in addition to a non-linear transformation task. Thus, in addition to evaluating multitasking, this framework captures the ability of the reservoir to balance two fundamental, but antagonistic properties: information storage and non-linear information processing [37, 39, 185]. The non-linear transformation task consists of regressing a slowly varying sinusoidal input signal to a different waveform, specifically, a square signal in this case [60, 163]. Performance is also measured using the ***R***^2^ coefficient of determination. Note that since the target signal is not time-lagged, this effectively consists in a one-step-ahead prediction, i.e., ***u***(*t*) is propagated to the input nodes at time *t* as per equation 6, but is not propagated within the reservoir; ***x***_*out*_(*t*) therefore only depends on (***u***(0), ***u***(1), …, ***u***(*t* − 1)).

A multitasking experiment, as performed using the synthetic hierarchical modular networks, proceeds as follows: The set of 4 modules encapsulated in the first of the 2 third-level modules is assigned memory capacity tasks. The other set is assigned non-linear transformation tasks (see Fig. 5). In the 2-task scenario, input nodes for each task correspond to all the nodes in a randomly selected first-level module. All the other first-level modules in the same third-level module are used as output. In the 4-task scenario, input nodes for each task also correspond to all the nodes in a first-level module. The other first-level module in the same second-level module is used as output. Finally, in the 8-task scenario, an input node is randomly selected in each of the 8 firstlevel modules. All of the other nodes in each module are used as output. When using more than 2 tasks, input signals are varied within a task category by using different seeds for the memory capacity task and varying the signal frequency for the non-linear transformation task. Specifically, the sinusoidal signal can complete either 10, 20, 30, or 40 complete cycles. Initial phase is also randomized. Note that the task assignments are maintained across networks of all three levels of the hierarchy.

As for the memory capacity task, we separately generated a training and a testing input signal of 2100 time points each and a warmup period of 100 time points was discarded from the resulting reservoir state trajectories. Different signals were also used for each network evaluated. All signals were propagated simultaneously across the network and a separate readout module was trained for each output module. The multitasking performance was assessed as the average ***R***^2^ score across all tasks.

### Hyperparameter tuning

Reservoir computing performance depends on many interacting hyperparameters, which are set a priori [37, 77, 147]. For example, spectral radius *α* can be tuned to control the reservoir dynamics and we study its effect on performance in the main analyses. Other features that pertain to the hierarchical modular network model are studied in the *Sensitivity analyses*. Finally, three hyperparameters are tuned via a random search [11] procedure: input gain *g*, pruning ratio *p*, and Tikhonov regularization parameter *λ*. The input gain scales the input matrix. The pruning ratio corresponds to the fraction of randomly selected edges to delete from the reservoir. Note that on average, this retains the relative density, degree, and modularity across hierarchical levels. Finally, the Tikhonov regularization parameter controls the strength of the L2 penalty in Ridge regression.

Hyperparameter tuning allows us to control for these parameters in subsequent analyses. It also contributes in addressing our research question by aggregating data across many experimental setups to obtain a high-performing regime that is not biased towards any topology or dynamical regime a priori. The procedure works as follows:

We start by sampling 125 hyperparameter configurations from uniform distributions over given ranges. Specifically, log_10_(*g*) ∼*U* (−10, 1), *p* ∼*U* (0, 1), and log_10_(*λ*) ∼*U* (−10, 1). In parallel, we consider random subsets of 25 synthetic networks from each hierarchical level and all 5 values of *α* considered in the main analyses. Then, using each hyperparameter configuration, we train and test each reservoir at the task at hand using the previously described procedures. The datasets are generated independently from the ones used to test the final model. Note that hyperparameter configurations are discarded if *p* does not retain strongly connected networks (any node must be reachable from any other node).

For each hyperparameter combination and network, we select the best performance across all spectral radii. We then average performance across all networks within each hierarchical level. Finally, we select the best performance across levels. The hyperparameter combination that leads to the best performance is retained. In this way, we obtain the hyperparameter regime with the best possible performance across dynamical regimes and hierarchical levels.

Note that for the empirical connectome and its null surrogates, we use the same procedure, but only tune input gain and regularization strength, maintaining the empirical topology. Furthermore, hyperparameter tuning is only performed using the empirical connectome, favoring it in the selected dynamical regime.

### Timescales

In line with previous definitions of neural timescales [24, 61, 126], we measure the timescale of a node’s activation timeseries as the time constant of an exponential decay function fit to the autocorrelation function (ACF):

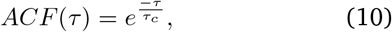

where *τ* is the time lag and *τ*_*c*_ is the characteristic timescale.

We calculate the autocorrelation function using the statsmodels open-source package (https://www.statsmodels.org/devel/) [166]. We only consider lags up to the first negative autocorrelation value in all timeseries considered. We then linearize the model by taking the logarithm of the resulting autocorrelation values and fit the characteristic time scale using least-squares fitting via numpy (https://numpy.org/citing-numpy/) [73]. Note that we only evaluate the output nodes’ timescales to avoid the biasing effect of the input signal. Furthermore, we discard the warmup period from the activation timeseries prior to estimating timescales.

### Graph analysis

#### Modularity maximization

To identify hierarchical modular partitions in the empirical connectome, we use the Louvain modularity maximization algorithm [18]. This algorithm detects non-overlapping communities of nodes that maximize the weight of within-community edges and minimize the weight of between-community edges using the quality function:

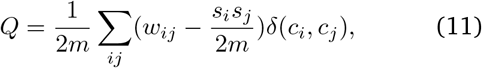

where *m* is the total weight of all edges in the network, *w*_*ij*_ is the weight of the edge incident on nodes *i* and *j, s*_*i*_ is the strength of node *i, c*_*i*_ is the community assignment of node *i* and *δ*(*c*_*i*_, *c*_*j*_) is the Kronecker delta function and is equal to 1 when *c*_*i*_ = *c*_*j*_ and 0 otherwise.

The algorithm recursively applies a sequence of two phases: modularity optimization, where nodal community labels are changed based on neighbouring labels, and community aggregation, where lower-level communities are nested into higher-order communities. Thus, the algorithm naturally yields hierarchical partitions. The algorithm was applied using the openly available *modularity_louvain_und* function from the Python version of the Brain Connectivity Toolbox (https://github.com/aestrivex/bctpy) [151]. To find a representative solution, we ran the algorithm 1000 times with different seeds and computed the z-scored Rand index [145] between every pair of first-level module partitions. We then chose the solution most similar to all others, retaining its higher-level modular partition. Note that we chose to select for representativeness at the finest resolution because the first-level modules are used to define reservoir inputs and outputs.

#### Clustering coefficient

For binary directed graphs such as the synthetic hierarchical modular networks, the clustering coefficient of a node corresponds to the fraction of all possible directed triangles around a node that exist. The clustering coefficient *C* of a node *u* can be defined as [52]:

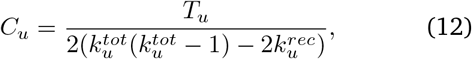

where *T*_*u*_ is the number of directed triangles through node *u*, 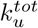 is the sum of its in- and out-degree, and 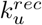 is its reciprocal degree.

It was computed using the openly available *clustering_coef_bd* function from the Python version of the Brain Connectivity Toolbox (https://github.com/aestrivex/bctpy) [151]. The network average clustering coefficient was computed as the mean across local clustering coefficients of all nodes in the network.

For weighted undirected graphs such as the empirical human connectome, the weighted local clustering coefficient of a node corresponds to the mean “intensity” of triangles around a node. The clustering coefficient *C* of a node *u* can be defined as [129]:

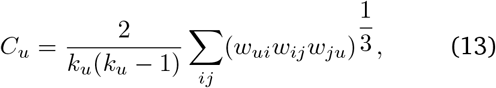

where *k*_*u*_ is the degree of node *u* and *w*_*ij*_ is the weight of the edge incident on nodes *i* and *j*, scaled by the largest weight in the network. It was computed using the openly available *clustering_coef_wu* function from the Python version of the Brain Connectivity Toolbox (https://github.com/aestrivex/bctpy) [151]. The module average clustering coefficient was computed as the mean across local clustering coefficients of all nodes in a module.

#### Cycles and motifs

A cycle is a path that begins and ends at the same node. A simple cycle additionally does not allow for repeated nodes. For the synthetic hierarchical modular networks, we compute all simple cycles up to length 5 using networkx [72], which implements a version of the algorithm of Gupta and Suzumura [69]. While we limit our analysis to short cycles, note that all networks considered have short diameter. Moreover, larger cycles are usually less relevant to network function and accounting for them is computationally prohibitive [53].

Motifs are local connection patterns that constitute the building blocks of a network’s functional repertoire [118]. Here, we study three-node motifs, for which there are 13 possible configurations. Specifically, we focus on functional motifs, which represent elementary information processing modes that can be engaged in a structural motif [171]. Functional motifs consist of all the subdivisions of a structural motif into the other structural motifs it contains. We count the number of each motif in a network using the Brain Connectivity Toolbox (https://sites.google.com/site/bctnet/) [151]. We then normalize these values by the total number of all motifs to obtain frequencies.

#### Communicability

Communicability between two nodes is defined as the length-weighted sum of all walks between them, with longer walks receiving progressively less weight [50]. The communicability matrix *C* of pairwise communicability estimates between all nodes in a network is calculated as the matrix exponential of the adjacency matrix *A*: *C* = *e*^*A*^. Following [35], we first normalize the adjacency matrix as *D*^1*/*2^*AD*^1*/*2^, where *D* is the diagonal weighted degree matrix. This normalization mitigates the disproportionate influence of high-strength nodes on communicability estimates.

### Data acquisition and connectome reconstruction

Magnetic resonance imaging (MRI) data from *n* = 327 unrelated, healthy young adults (28.6 ± 3.73 years old, 55% female) from the Human Connectome Project (HCP) were used to reconstruct structural connectivity networks and build a group-representative connectome [10, 158]. Participants were scanned at Washington University using the HCP’s customized 3-Tesla Siemens “Connectome Skyra” MRI scanner. The protocol included (1) a magnetization-prepared rapid acquisition gradient echo (MPRAGE) sequence (TR = 2400 ms, TE = 2.14 ms, FOV = 224 mm × 224 mm, voxel size = 0.7 mm^3^, 256 slices) and (2) a spin-echo echo-planar imaging (EPI) sequence (TR = 5520 ms, TE = 89.5 ms, FOV = 210 mm ×180 mm, voxel size = 1.25 mm^3^, b-value = 1000, 2000, and 3000 s/mm^2^, 270 diffusion directions, 18 b0 volumes). All participants provided informed written consent and the protocol was approved by the Washington University Institutional Review Board. Further details on data acquisition can be found in [184].

The HCP minimal preprocessing pipelines [65] were applied to the MRI data and streamline tractography tools from the MRtrix3 open-source software [181] were used to reconstruct structural connectivity networks in individual participants from diffusion-weighted MRI data. The MPRAGE volume was segmented into white matter, grey matter, and cerebrospinal fluid to perform anatomically constrained tractography. Grey matter was parcellated into 400 regions according to the Schaefer functional atlas [155]. Fiber orientation distributions were generated using the multi-shell multi-tissue constrained spherical deconvolution algorithm from MR-trix3 [29, 86]. Tractograms were initialized with 40 million streamlines and constrained with a maximum tract length of 250 and a fractional anisotropy cutoff of 0.06. A spherical deconvolution-informed filtering procedure (SIFT2) was then applied following Smith et al. [167] to estimate streamline-wise cross-section multipliers. For additional details on MRI data preprocessing and connectome reconstruction, see [134].

Group-representative structural networks were then generated to amplify signal-to-noise ratio using functions from the netneurotools open-source package (https://netneurotools.readthedocs.io/en/latest/index.html). A consensus approach was adopted to preserve (1) the mean density across participants and (2) the participant-level edge length distribution [16]. First, the cumulative edge length distribution across individual structural connectivity matrices is divided into *M* bins, with *M* corresponding to the average number of edges across participants. The edge occurring most frequently across participants is then selected within each bin, breaking ties by selecting the edge with the highest average weight. This procedure is performed separately for intra- and inter-hemispheric edges to ensure that the latter are not under-represented. The resulting edge set constitutes the distance-dependent group-consensus structural network. The weight of each edge is then computed as the mean across participants. Finally, the weights were logtransformed and rescaled to the [0, 1] range to reduce variability.

### Intrinsic timescales from magnetoencephalography (MEG)

The *neuromaps* (https://github.com/netneurolab/neuromaps) [110] toolbox was used to obtain an intrinsic timescale map derived from MEG. The map was downloaded in its native fsLR4k space. The Schaefer-400 atlas in fsLR32k space was then downsampled using nearest neighbour interpolation to parcellate the map in 400 cortical regions. Details regarding data acquisition and processing are available at [157]. Briefly, resting-state MEG data were acquired in a subset of n = 33 unrelated healthy young adults (22-35 years old, 16 female) from the Human Connectome Project (HCP). The 6-minute scans were sampled at 2034.5 Hz and processed using Brainstorm [175]. MEG recordings were co-registered with individual MRI scans, downsampled to 509 Hz, and preprocessed using notch filters at 60, 120, 180, 240, and 300 Hz, along with a high-pass filter at 0.3 Hz to remove slow-wave and DC-offset artifacts. Artifacts such as heartbeats, eye movements, and muscle activity were removed using automatic procedures involving electrocardiogram (ECG) and electrooculogram (EOG) recordings and Signal-Space Projection (SSP). Source reconstruction was performed using linearly constrained minimum variance (LCMV) beamforming on the fsLR4k cortical surface, with data covariance regularization and noise covariance normalization. Finally, intrinsic neural timescales were derived from source-level power spectral densities using the FOOOF toolbox [46], which decomposes power spectra into periodic and aperiodic components. The “knee frequency” from the aperiodic fit was used to compute the intrinsic timescales over the frequency range of 1-60 Hz.

### Network null models

#### Degree-preserving rewiring

To randomize network topology while preserving network size (number of nodes), density (proportion of present edges), and degree sequence (number of edges incident to each node), we applied the classic switching method [111] using the openly available *randmio_dir_connected* function from the Python version of the Brain Connectivity Toolbox (https://github.com/aestrivex/bctpy) [151]. The method was applied to the third-level hierarchical modular networks to generate an ensemble of degree-preserving randomized surrogates.

#### Hierarchical modular null model

To randomize a network while preserving its (hierarchical) modular architecture in addition to its size, density, and degree sequence, we develop a novel network null model building on conventional degree-preserving rewiring. Our model draws on an existing modular random graph model [153] and the intuitive notion that modularity can be operationalized as the relative density of intra-modular connections [59, 127]. For a strictly modular network null model, separately rewiring within and between-module edges suffices to randomize a network while preserving intra-modular density [153]. Generalizing to hierarchical modularity, it can be inferred that *L* nested partitions will result in *L* + 1 subgraphs of fixed density in order for each module of each partition level to maintain intra-modular connectivity (density within-module at level *l* + 1 but between modules at level *l* must also be maintained). The algorithm implementing this model proceeds as follows:

Consider *G*_*e*_ = (*V, E*_*e*_), the empirical graph to be randomized with vertex set *V* and edge set *E*_*e*_ and *G*_*c*_ = *K*_*n*_ = (*V, E*_*c*_), the complete graph of the same size *n* = |*V* |, where *E*_*c*_ contains all possible edges. Next, consider the series of nested partitions 𝒫^(1)^, 𝒫^(2)^, …, 𝒫^(*L*)^, where 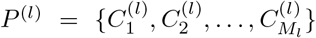 is the partition at level *l*, 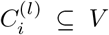 is community *i* at level *l* and *M*_*l*_ is the number of communities at level *l*.

We start by creating hierarchical decompositions of *G*_*e*_ and *G*_*c*_ as follows:

1. We create *l* subgraphs of *G*_*c*_ that consist of the complete graphs of all communities at level *l*, respectively:

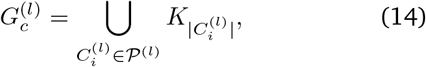 The final level is the fully connected graph 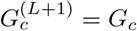
2. We then build the corresponding empirical subgraphs:

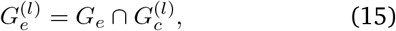

as the intersections of *G*_*e*_ and 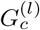
3. For both *G*_*e*_ and *G*_*c*_, we then build the incremental subgraphs:

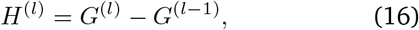

for *l* = 2, …, *L* + 1. Subgraphs *H*^(*l*)^ are therefore difference-based subgraphs corresponding to the additional connections introduced at level *l*, with *H*^(1)^ = *G*^(1)^ as the base case.
4. Next, for each empirical incremental subgraph 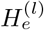, we apply degree-preserving rewiring [111], while introducing the additional constraint that the edge swap:

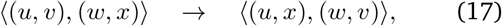

is only accepted if (*u, x*) and (*w, v*) also exist in 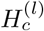. This procedure results in the rewired incremental subgraphs 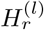.
5. Finally, the hierarchical modularity-preserving randomized network *G*_*r*_ is obtained by summing the rewired incremental subgraphs 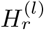:

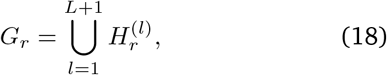

## ACKNOWLEDGMENTS

We thank Justine Hansen, Eric Ceballos, Vincent Bazinet, Zhen-Qi Liu, Asa Farahani, Yigu Zhou, Tahmineh Taheri, Moohebat Pourmajidian, and Aleksandar Mihajlovski for helpful comments. F.M. acknowledges support from the Fonds de Recherche du Québec—Nature et Technologies, the Healthy Brains for Healthy Lives initiative and the Centre Union Neurosciences and Artificial Intelligence—Quebec. B.M. acknowledges support from the Natural Sciences and Engineering Research Council of Canada, Canadian Institutes of Health Research, Brain Canada Foundation Future Leaders Fund, the Canada Research Chairs Program, the Michael J. Fox Foundation and the Healthy Brains for Healthy Lives initiative.

---

**Figure S1.**
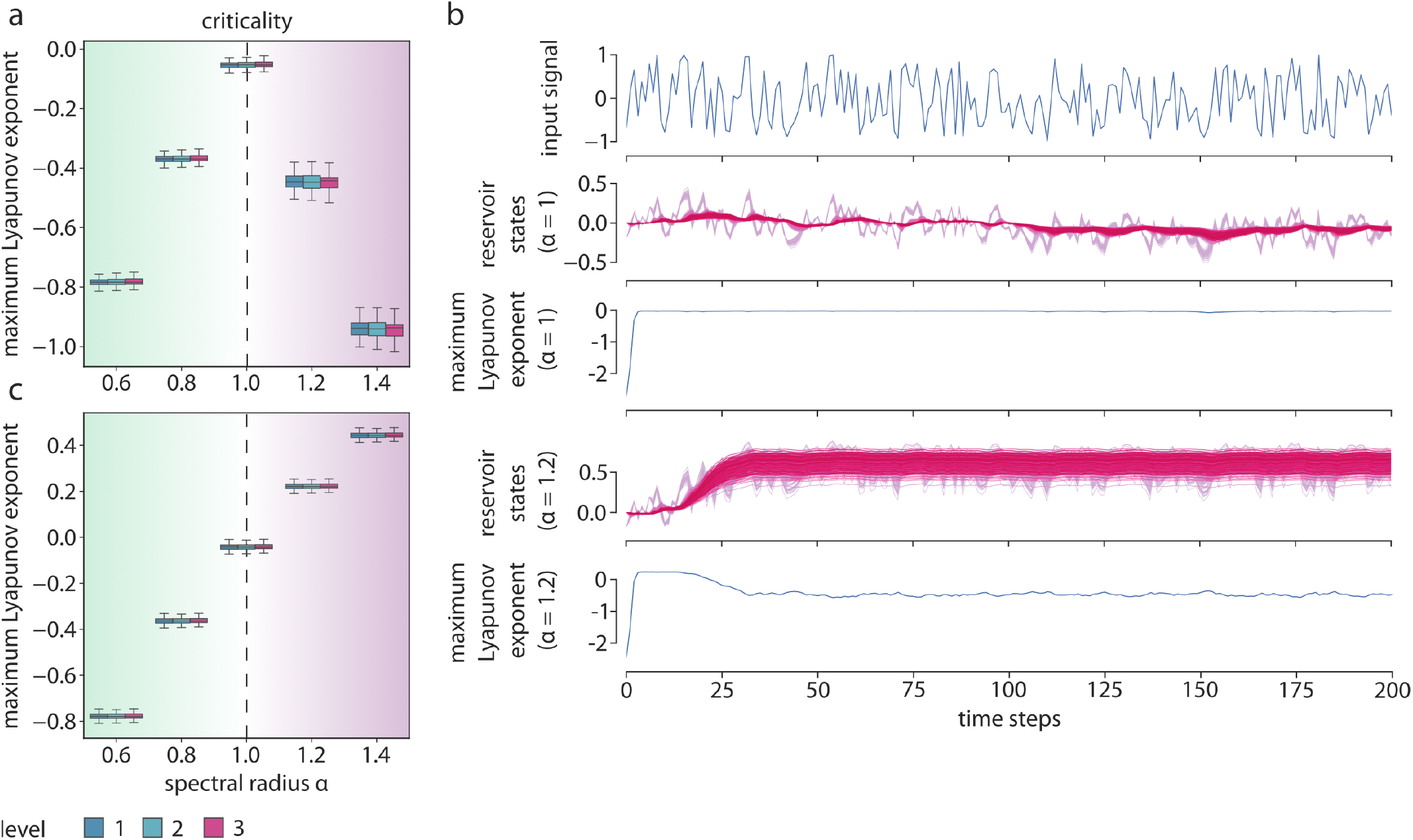
Lyapunov exponent analysis. (a) Maximum Lyapunov exponent (LE) as a function of the spectral radius *α* across reservoirs of tanh units for each hierarchical level. Criticality is characterized by a negative maximum LE close to zero. Beyond *α* = 1, maximum LE decreases instead of crossing to chaos (positive LE) because of tanh unit saturation. (b) From top to bottom: Example input signal for the memory capacity task. Reservoir states at criticality for an example third-level network and the corresponding instantaneous contributions to the maximum Lyapunov exponent (obtained using a QR decomposition algorithm). Reservoir states for the same network at *α* = 1.2 and the corresponding instantaneous contributions to the maximum Lyapunov exponent (notice the positive values prior to saturation). Different colours correspond to different nodes. (c) Maximum Lyapunov exponent (LE) as a function of the spectral radius *α* across linear reservoirs for each hierarchical level. Comparatively to the tanh units, linear units do not saturate, leading to positive maximum LEs consistent with chaos for *α >* 1.

**Figure S2.**
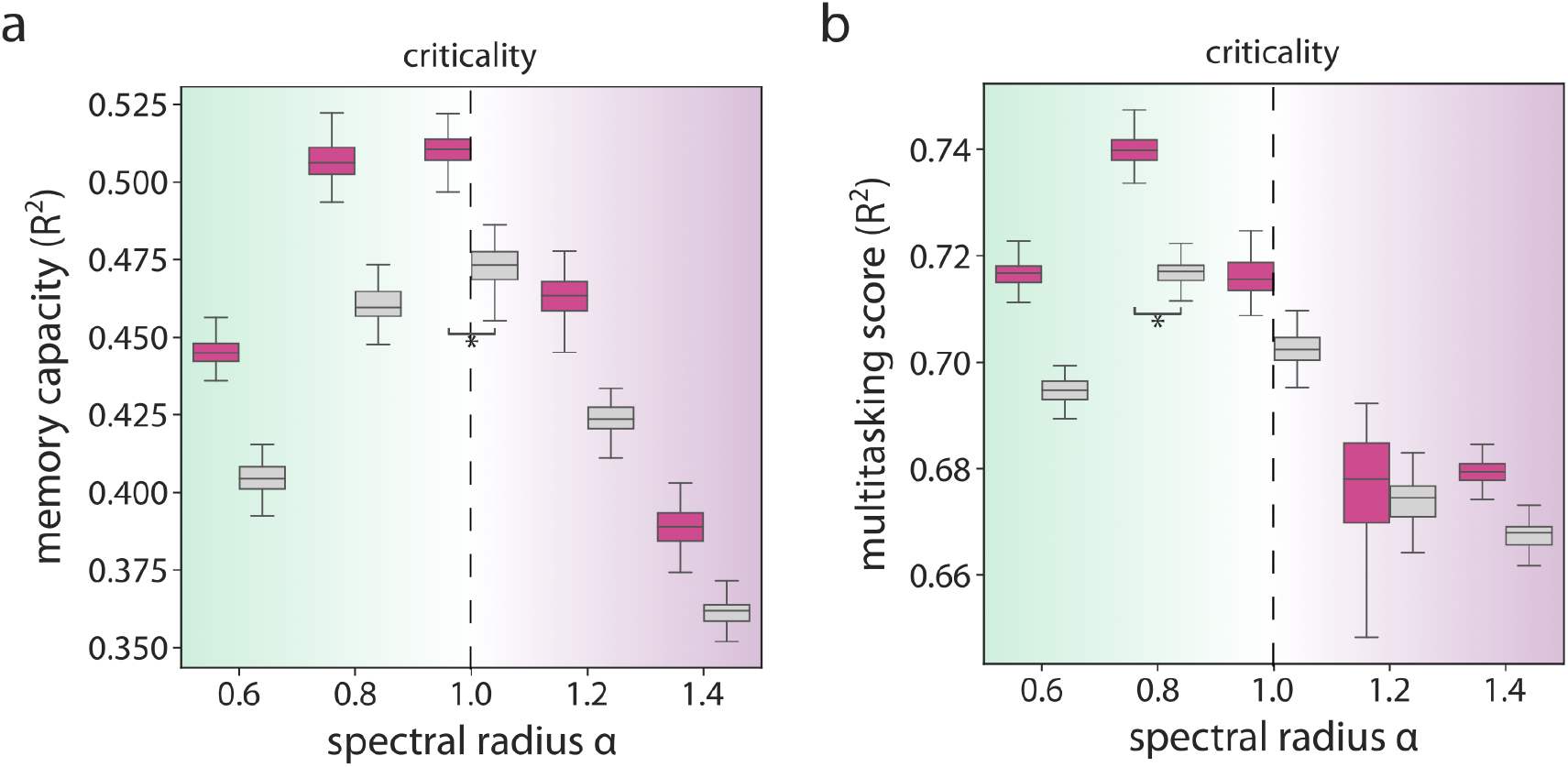
Contrast between hierarchical modular and random reservoir performance. Memory capacity (a) and multitasking score (b) as a function of the *α* parameter for level 3 networks (magenta) and their degree-preserving randomized surrogates (grey). Asterisks indicate statistical significance of differences between levels according to Wilcoxon-Mann-Whitney tests (*p <* 10^−33^). Significance is only shown for the highest performing regime.

**Figure S3.**
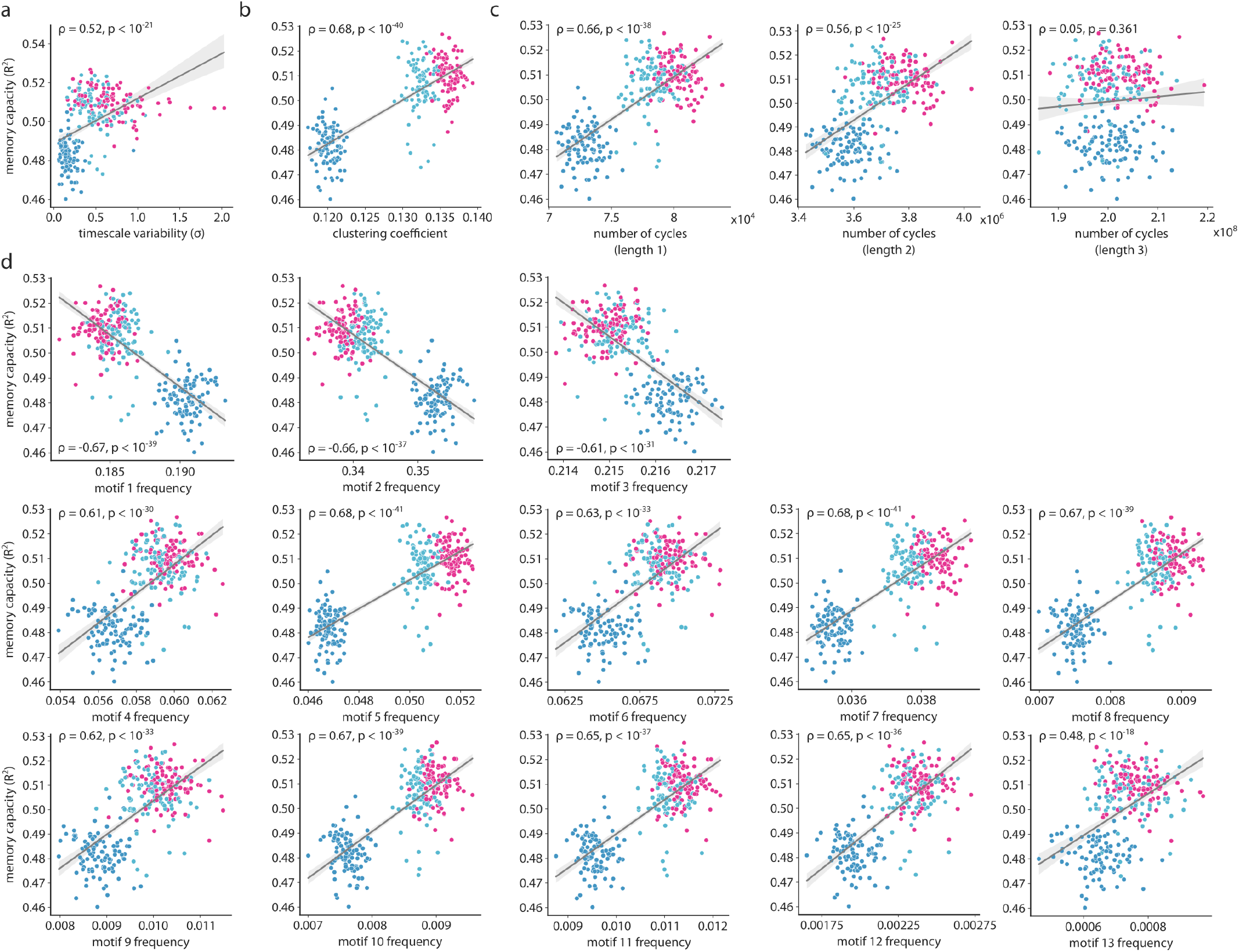
Determinants of (hierarchical) modular reservoir performance. Relationship between memory capacity (*R*^2^) of synthetic (hierarchical) modular reservoirs and their timescale variability (a; in units of standard deviation), clustering coefficient (b), number of cycles of different length (c), and motif frequency (d). Each point represents a network. Linear regression lines (grey) and 95% bootstrapped confidence intervals (shaded bands; 1000 samples) are shown for visualization purposes. Spearman correlation coefficients (*ρ*) and associated p-values are shown in each scatter plot. Level 1 networks are shown in blue, level 2 networks are shown in teal, and level 3 networks are shown in magenta.

**Figure S4.**
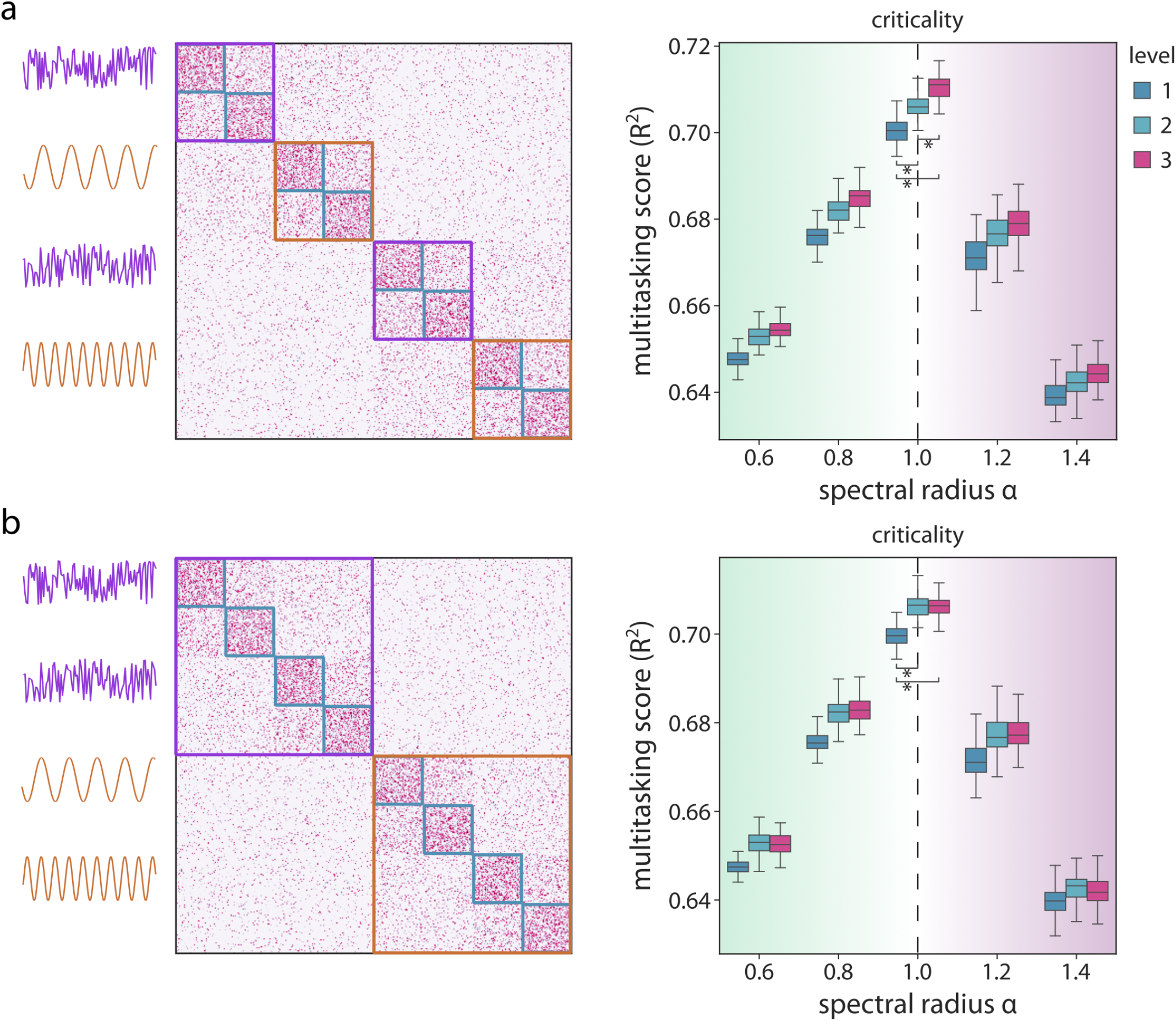
Hierarchical modularity supports multitasking - 4. tasks. (a) Left: Schematic of the multitasking framework: the 4 second-level modules are each assigned a different task. The memory capacity task and the non-linear transformation task are interleaved. Both input signals are propagated simultaneously across the network. Right: Average *R*^2^ regression score across tasks as a function of the *α* parameter for each hierarchical level. (b) Same as (a) when task categories are not interleaved (the 2 third-level modules are each assigned a different task category). Asterisks indicate statistical significance of differences between levels according to Wilcoxon-Mann-Whitney tests (*p <* 10^−17^). Significance is only shown for the highest performing regime.

**Figure S5.**
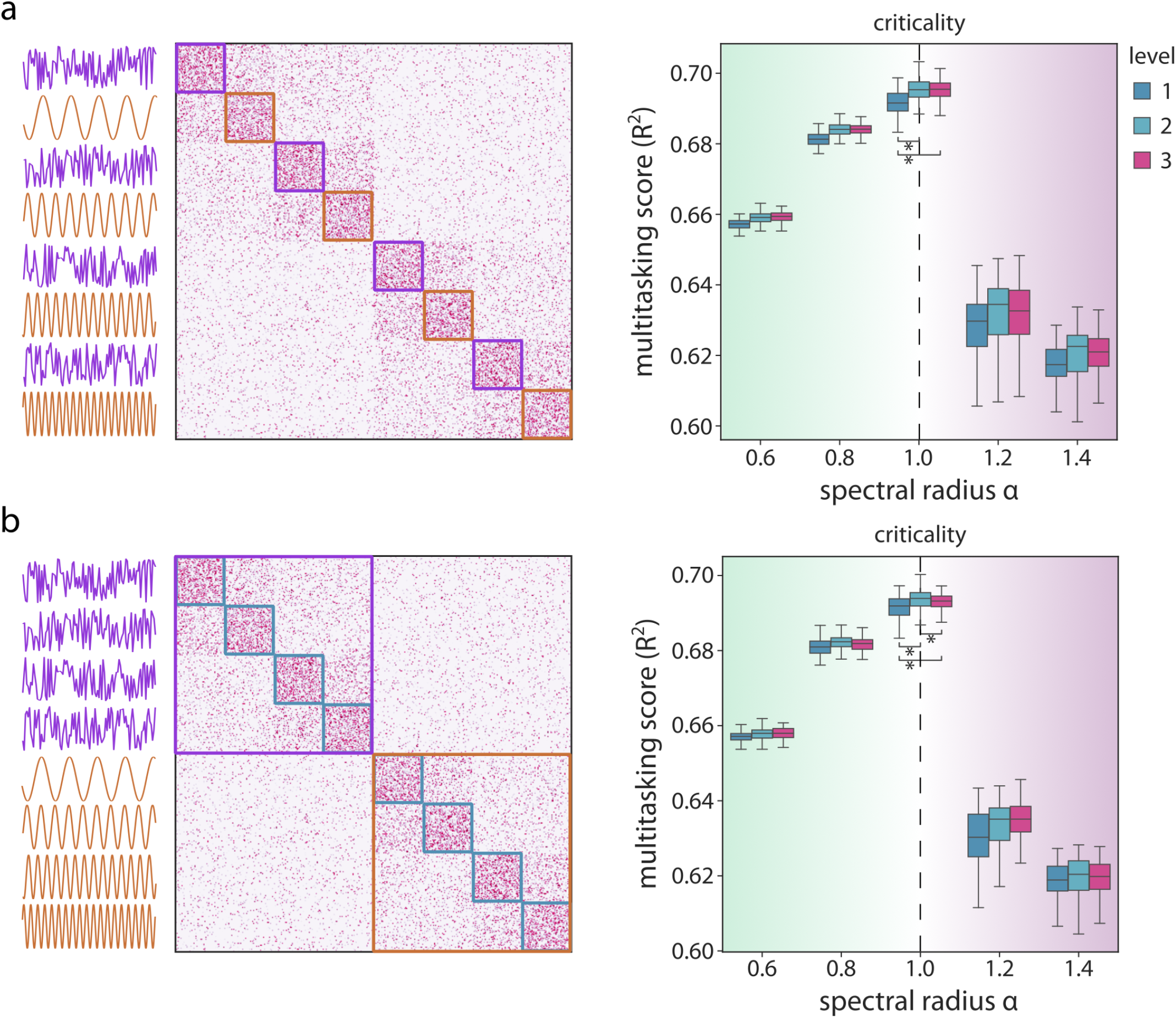
Hierarchical modularity supports multitasking - 8. tasks. (a) Left: Schematic of the multitasking framework: the 8 first-level modules are each assigned a different task. The memory capacity task and the non-linear transformation task are interleaved. Both input signals are propagated simultaneously across the network. Right: Average *R*^2^ regression score across tasks as a function of the *α* parameter for each hierarchical level. (b) Same as (a) when task categories are not interleaved (the 2 third-level modules are each assigned a different task category). Asterisks indicate statistical significance of differences between levels according to Wilcoxon-Mann-Whitney tests (*p <* 0.05). Significance is only shown for the highest performing regime.

**Figure S6.**
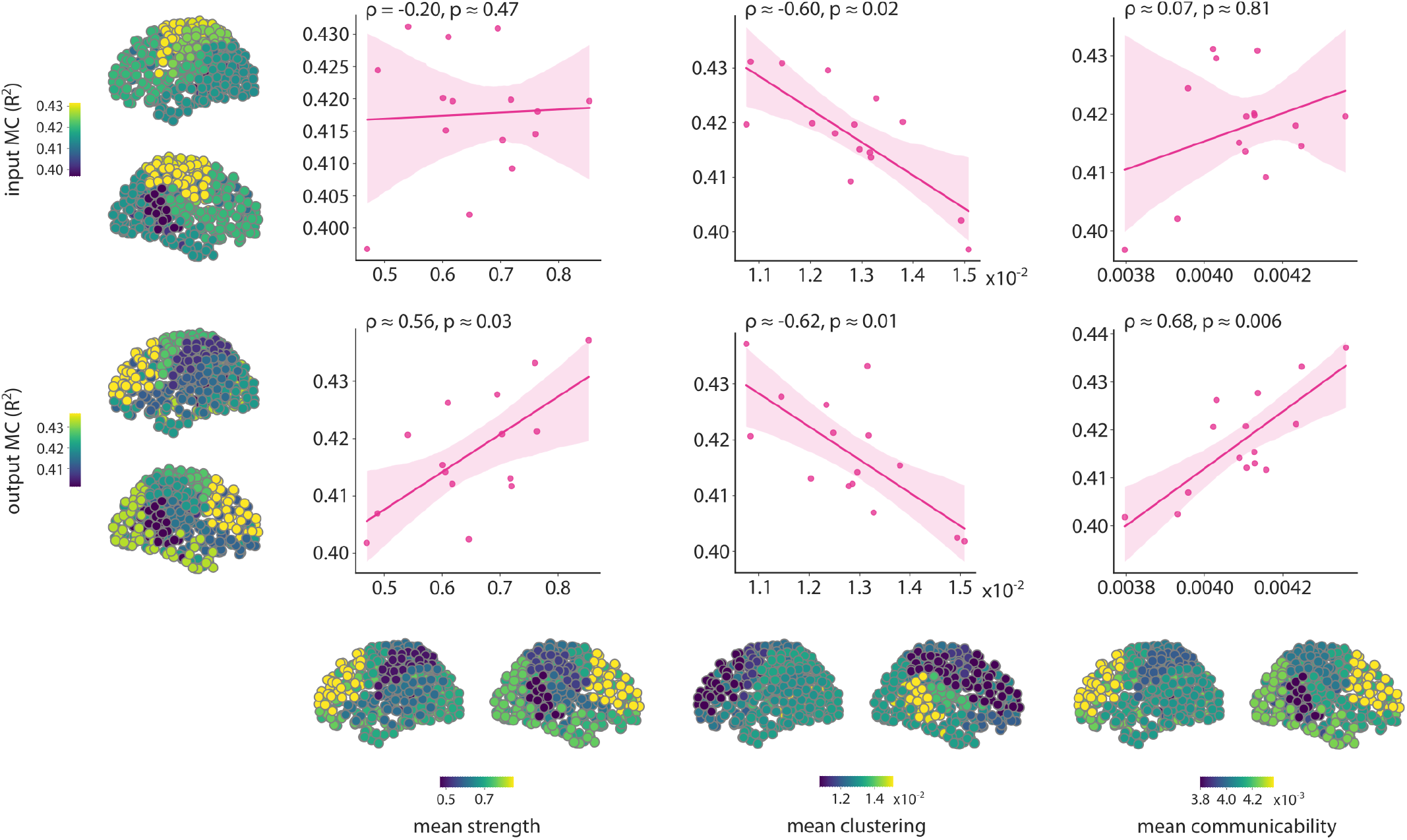
Network determinants of connectome module performance. Relationships between module-wise memory capacity (MC; *R*^2^) as input (top) and output (bottom) as a function of module-average strength (left), clustering (middle), and communicability (right). Each point represents a module. Linear regression lines (magenta) and 95% bootstrapped confidence intervals (shaded bands; 1000 samples) are shown for visualization purposes. Spearman correlation coefficients (*ρ*) and associated p-values are shown on top of each scatter plot.

**Figure S7.**
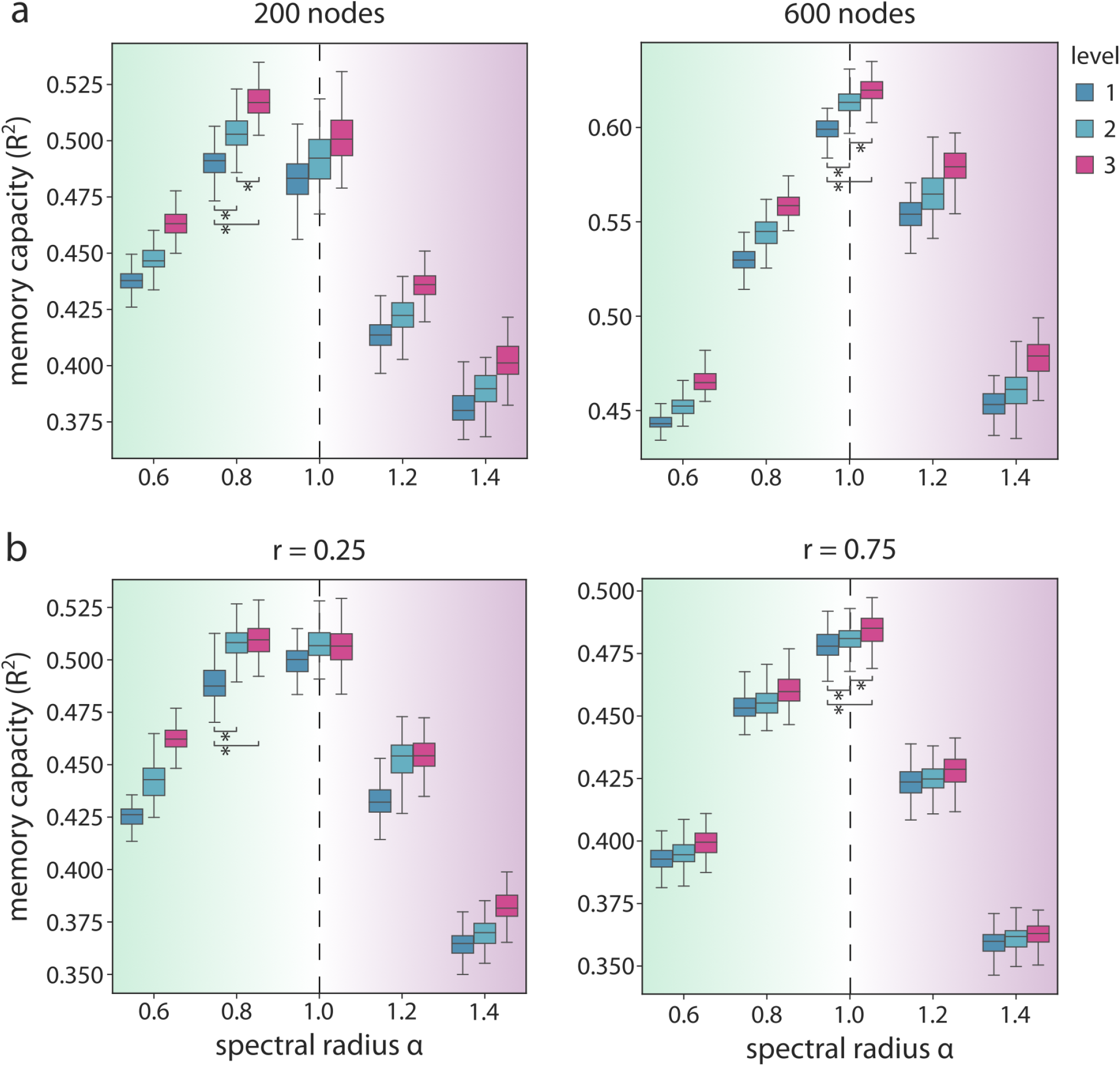
Sensitivity analyses - network size and edge probability scaling. Memory capacity (*R*^2^) as a function of the *α* parameter across hierarchical levels, for different network sizes (a; 200 and 600 nodes) and different edge probability scaling factors *r* (b; 0.25 and 0.75). Asterisks indicate statistical significance of differences between levels according to Wilcoxon-Mann-Whitney tests (*p <* 0.01). Significance is only shown for the highest performing regime.

**Figure S8.**
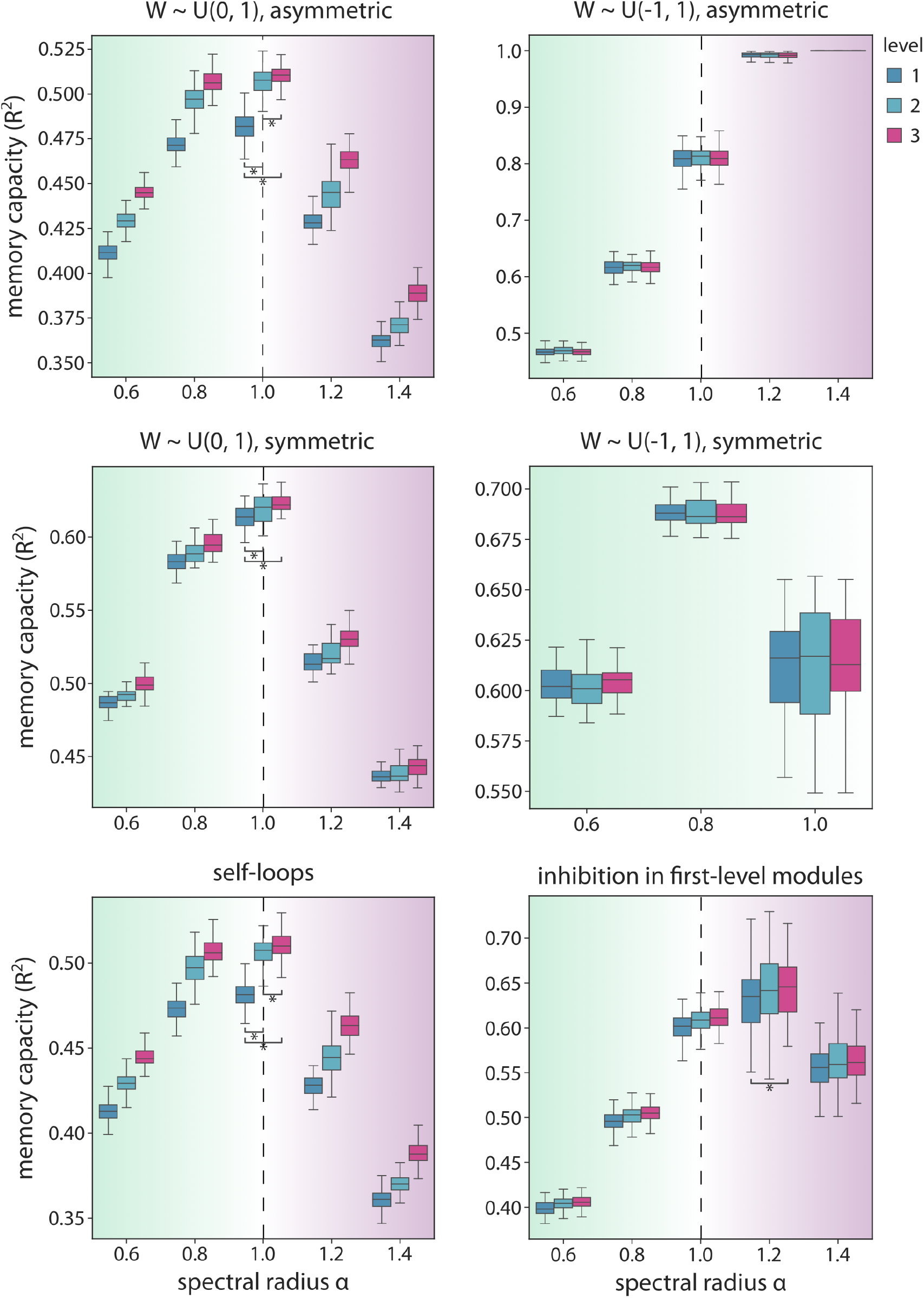
Sensitivity analyses - weight distribution, symmetry, and self-loops. Memory capacity (*R*^2^) as a function of the *α* parameter across hierarchical levels, for asymmetric positive networks (top, left), asymmetric signed networks (top, right), symmetric positive networks (middle, left), symmetric signed networks (middle, right), asymmetric positive networks containing self-loops (bottom, left), and asymmetric signed networks only containing negative weights in the first-level modules (bottom, right). Asterisks indicate statistical significance of differences between levels according to Wilcoxon-Mann-Whitney tests (*p <* 0.05). Significance is only shown for the highest performing regime. Note that we only show the stable regime for symmetric signed networks for visualization purposes as performance considerably drops in the chaotic regime.

1 Note that our matrices are also positive, ensuring diagonal Schur stability for *α <* 1 [196]. However, in practice, tuning the spectral radius below 1 usually suffices to obtain the echo state property for any matrix [103].

